# Macrophage migration-dependent retraction fibers and migrasomes - a structural basis for spatiotemporal cytokine secretion

**DOI:** 10.1101/2024.12.12.627632

**Authors:** Xichun Li, Azadeh Anbarlou, Vrushali Maste, Nicholas D. Condon, Darren Brown, Hongyu Shen, Hao-Ruei Hsu, Amina Ashraf, Sylvia Tan Jinhui, Wanyi Wang, Oleksandr Chernyavskiy, Richard M. Lucas, Lin Luo, Yvette W. H. Koh, Rachael Z. Murray, Graham J. Lieschke, Jennifer L. Stow

**Author notes:** Corresponding authors: Prof. Jennifer L. Stow, Institute for Molecular Bioscience, The University of Queensland, 306 Carmody Road, St Lucia Brisbane QLD 4072 Australia, E, Prof. Graham J. Lieschke, Australian Regenerative Medicine Institute, 15 Innovation Walk, Monash University, Clayton VIC 3800 Australia. E. These authors contributed equally (Xichun Li, Azadeh Anbarlou, Graham J. Lieschke, Jennifer L. Stow).

## Abstract

Migrating cells leave behind trails of matrix-bound retraction fibers and migrasomes, but their cell-specific physiological functions are largely unknown. We comprehensively characterize retraction fibers and migrasomes of migrating macrophages in mouse and zebrafish *in vitro* and *in vivo* models. These structures share common and macrophage-centric components of tetraspanin enriched microdomains (TEMs) including tetraspanin 4 (TSPAN4), the integrin CD11b and the pTRAP, SCIMP; like TSPAN4, we show that SCIMP expression modulates retraction fiber and migrasome formation. In immune-activated migrating macrophages, rear-positioned recycling endosomes regulate the secretory trafficking of newly synthesized inflammatory cytokines into retraction fibers and migrasomes. Transmembrane TNF is delivered via SNARE-mediated carriers to the cell surface and to the surfaces of both retraction fibers and migrasomes, from where it can be released. Retraction fibers are thus newly identified here as cytokine secretion sites, and along with migrasomes they allow migrating macrophages to secrete more cytokine compared to stationary cells. Whereas cytokine secretion has been viewed as a process of *ad hoc* diffusion, instead, we reveal a new mechanism of migration-dependent, localized deposition and spatiotemporal release of bioactive cytokines from RFs and migrasomes. Cytokine deployment is thus organized in a constrained manner facilitating direct influence on local inflammatory responses.

## INTRODUCTION

Macrophages play a central role in innate immunity, serving to surveil tissues for the detection and removal of pathogens and to respond to injury (1, 2). Along with other innate immune cells, macrophages are actively recruited as early cellular responders, migrating to and at sites of wounding or infection (3–5). Macrophage activation leads to responses that include the phagocytosis of pathogens and debris, and the temporal release of proinflammatory mediators, cytokines and chemokines, aimed at sterilizing sites and warding off infection, followed later by release of anti-inflammatory mediators that promote resolution of inflammation and tissue repair (2, 6). Despite the inherently migratory nature of activated macrophages responding to threats *in vivo,* paradoxically much of our current understanding of macrophage inflammatory outputs derives from the analysis of non-migratory macrophages already at sites of infection. Even more of this knowledge is based on *in vitro* observations of stationary macrophages under culture conditions. Hence macrophage inflammatory responses mounted during the movement of these migrating cells remain poorly understood.

Recent studies on cell migration have rekindled interest in the presence of membranous retraction fibers (RFs) that emerge from the rear of migrating cells and, although seen much earlier (7), newer studies largely from the group of Li Yu (8, 9) have documented and characterized the extensive trails of RFs that bind to matrix substrates and are left behind by migrating fibroblasts and other cells in culture. Most strikingly they found that large, spherical ‘migrasomes’ formed on retraction fibers are a new source of extracellular vesicles (8). The RFs and migrasomes are replete with tetraspanin molecules clustered in tetraspanin-enriched microdomains (TEMs) which pack together into higher-order structures that swell to form migrasomes at branch points along RFs (10). These TEMs also include concentrated sphingolipids and cholesterol, as well as the integrins that attach the migrasomes and RFs to the extracellular matrix (10–12). Migrasomes have been ascribed several physiological roles, such as extruding damaged mitochondria (13) and releasing chemokines and growth factors in model organisms (14–16) and in cultured fibroblasts (17). While up to now migrasomes have been emphasized as the main perpetrators of these functions, RFs themselves share many of the same components and they give rise to vesicular ‘retractosomes’ although no function has yet been ascribed to these vesicles (18).

Advancing an understanding of RFs and migrasomes now requires detailed investigation of their production and functions in a cell type specific manner in complementary *in vivo* and *in vitro* settings. The current study achieves this by examining the composition and fates of RFs and migrasomes in migrating macrophages which are analyzed during *in vivo* migration in a zebrafish model and *in vitro* in cultured mouse primary macrophages and a cell line.

The outcomes reported here reveal new macrophage-centric inflammatory regulators as components of migrasomes and RFs, including the canonical macrophage integrin and inflammatory mediator CD11b (19) and a TEM component, the pTRAP signaling adaptor and proinflammatory regulator SCIMP (20–22). The formation and degradation of macrophage RFs and migrasomes are documented in *in vitro* and *in vivo* contexts. A pivotal role of immune-activated macrophages is their secretion of proinflammatory cytokines (23, 24) and here we show that cytokines, including tumor necrosis factor alpha (TNF), are released during macrophage migration. Indeed the secretion of TNF is enhanced and spatiotemporally defined by the deployment of RFs and migrasomes as vehicles for the distribution and secretion of this clinically-significant cytokine. These findings highlight the need to reimagine and redefine inflammatory responses perpetrated specifically by migrating cells, where distinctive cell morphologies, rear-polarized intracellular trafficking and the extrusion of significant swathes of membrane, can influence inflammatory outputs in tissue niches.

## RESULTS

### Migrating macrophages leave persistent trails of retraction fibers (RFs) and migrasomes

Mouse primary bone marrow derived macrophages (BMMs), cells of the RAW264.7 macrophage cell line and human monocyte-derived macrophages, plated on fibronectin (FN) and stimulated with CSF-1, undergo random migration leaving behind elaborate membrane trails of long, thin RFs with attached spherical migrasomes, both stained with fluorescent wheat germ agglutinin (WGA) (**Fig 1A and S1A Fig**). These migration-dependent RFs and migrasomes generated *in vitro* are similar in appearance to those reported in cultured fibroblasts and other cell types (8, 9). We also examined zebrafish macrophages migrating towards a tail wound site *in vivo*, typically captured in a 3-day post-fertilization embryo by confocal microscopy (**Fig 1B** and **Video 1)**. Using a macrophage reporter line Tg(mfap4:mTurquoise2;mpeg1:mCherry-CAAX) that is biologically neutral with respect to RF and migrasome formation, higher resolution live confocal imaging shows RFs emerging from the rear of migrating cells and trails of migrasome-like structures *in vivo* (**Fig 1C, insets**). Using live lattice light sheet microscopy (LLSM), straight or branched/forked RF configurations were visible *in vivo* (**Fig 1D**) and at this spatiotemporal resolution, 1-3 RFs per cell were detected in 34% of randomly-selected zebrafish macrophages (**Fig 1E**). These RFs were up to 60 μm long (**Fig 1F**) and visible attached to the migrating cells for up to 270 sec (**S1B Fig**). Use of another biologically neutral cytoplasmic reporter, Tg(mpeg1:Gal4FF;UAS:NTR-mCherry) confirms that the macrophage RFs and migrasomes (**Figs 1G** and **H**) carry cytoplasmic material *in vivo* (**Figs 1G, H** and **Video 2**). The confocal movie panels in **Fig 1H** show a long RF with internal cytoplasm and attached migrasomes at the distal end (arrows). Furthermore, FACS-isolated primary zebrafish macrophages migrating *in vitro* on FN-coated plates also display elongated WGA-staining processes consistent with RFs (**S1C Fig**, left image) beaded with migrasome-like swellings (**S1C Fig**, right image). From these data collectively, we conclude that RFs and migrasomes are a prominent feature of macrophage migratory behavior *in vivo* and *in vitro*.

**Figure 1.**
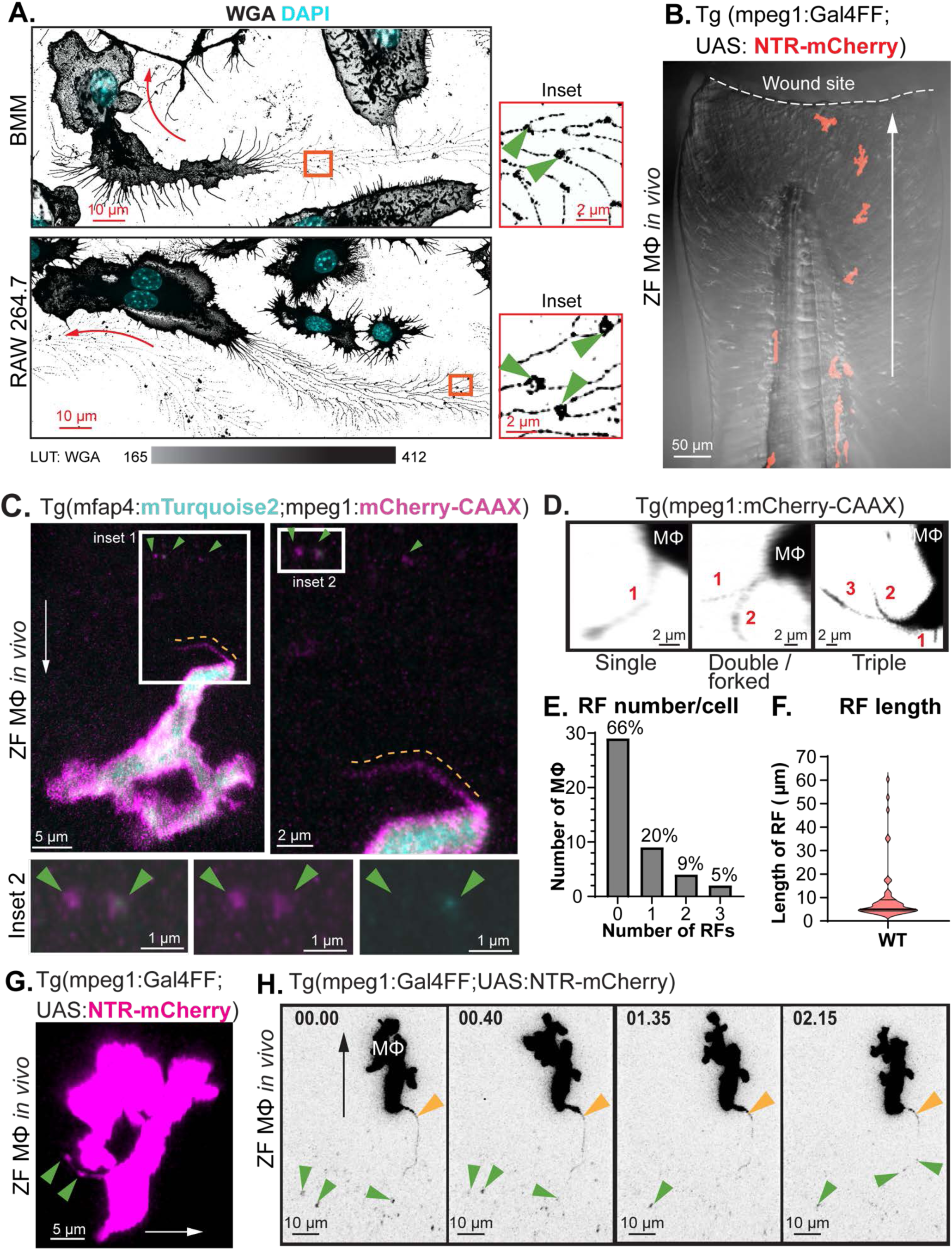
Retraction fibers and migrasomes of migrating macrophages *in vitro* and *in vivo*. **A.** Mouse bone marrow-derived macrophages (BMMs) and RAW264.7 cells on FN-coated dishes, in CSF-1 for 16 hrs were then fixed and stained with Alexa Fluor 488-WGA and DAPI. Green arrowheads indicate migrasomes; red arrows indicate cell migration direction. LUT: Alexa Fluor 488-WGA, inverted Grays; DAPI, inverted Cyan. **B.** Macrophages migrating to a tail transection wound site in a zebrafish embryo *in vivo*. Tg(mpeg1:Gal4FF;UAS:NTR-mCherry) macrophages imaged by confocal microscopy (red) are superimposed on a bright field image of the tail and wound site (dashed line). White arrow indicates migration direction. See **Video 1**. **C-G.** RF and migrasome formation by zebrafish macrophages *in vivo* in two reporter lines. **C-E** from Tg(mfap4:mTurquoise2;mpeg1:mCherry-CAAX) macrophages. **C.** Macrophage with RFs and migrasomes. Right panel is inset 1 detail, and right bottom panels are inset 2 details. Orange dashed lines indicate RFs; green arrowheads indicate migrasomes. White arrow indicates migration direction, determined from timelapse video context. **D.** Examples of single, double and triple RF configurations, images on LLSM Zeiss LLS7. Red numbers indicate RFs. **E.** Bar graph shows frequency of RF configurations in single cells. Measurement was performed on confocal live imaging. n = 44 cells. **F.** Distribution of RF lengths (n=60 RFs). **G-H** are from Tg(mpeg1:Gal4FF;UAS:NTR-mCherry) macrophages**. G.** Trailing migrasomes (green arrowheads). **H.** Time series showing RF (orange arrowheads) and trailing migrasomes (green arrowheads). See **Video 2**. **A-H**, scalebars as indicated.

Tetraspanin-enriched microdomains (TEMs) are structural components of RFs and migrasomes and the tetraspanin, TSPAN4-EGFP has consistently been expressed in cells as a label for these structures (10, 11). In transiently-transfected, migrating RAW264.7 macrophages, the RFs and migrasomes colabel with WGA and TSPAN4-EGFP (**Fig 2A**). Overexpression of TSPAN4-EGFP in these cells results in exaggerated RF trails (**Fig 2A**) reflecting earlier observation of this expansion in other cell types (10, 18). In live zebrafish macrophages *in vivo*, selective expression of a tspan4a-EGFP reporter shows widespread tspan4a-EGFP in the soma, overlapping the cytoplasmic NTR-mCherry marker (**Fig 2B** and **Video 3**). A representative movie depicts a macrophage with two RFs marked by the tspan4a-EGFP reporter as they change shape and mature through stages of RF beading and the appearance of migrasomes. Of note, in fish, the tspan4a-EGFP reporter is not biologically neutral, significantly increasing the number of migrasomes per macrophage without a detectable increase in their size *in vivo* (**Figs 2C** and **D**) and increasing RF numbers from 2 up to 5 per cell (**S2A** and **S2B Fig**). A normalized parameter based on RF number and duration per cell confirms a significant increase in tspan4a-EGFP RFs (**Fig 2E**). Finally, the ZF4 zebrafish fibroblast cell line forms RFs and migrasomes *in vitro* (**S2C Fig**), with WGA-stained, donut-shaped migrasomes frequently reported by others in fibroblast cultures (8, 10, 11).

**Figure 2.**
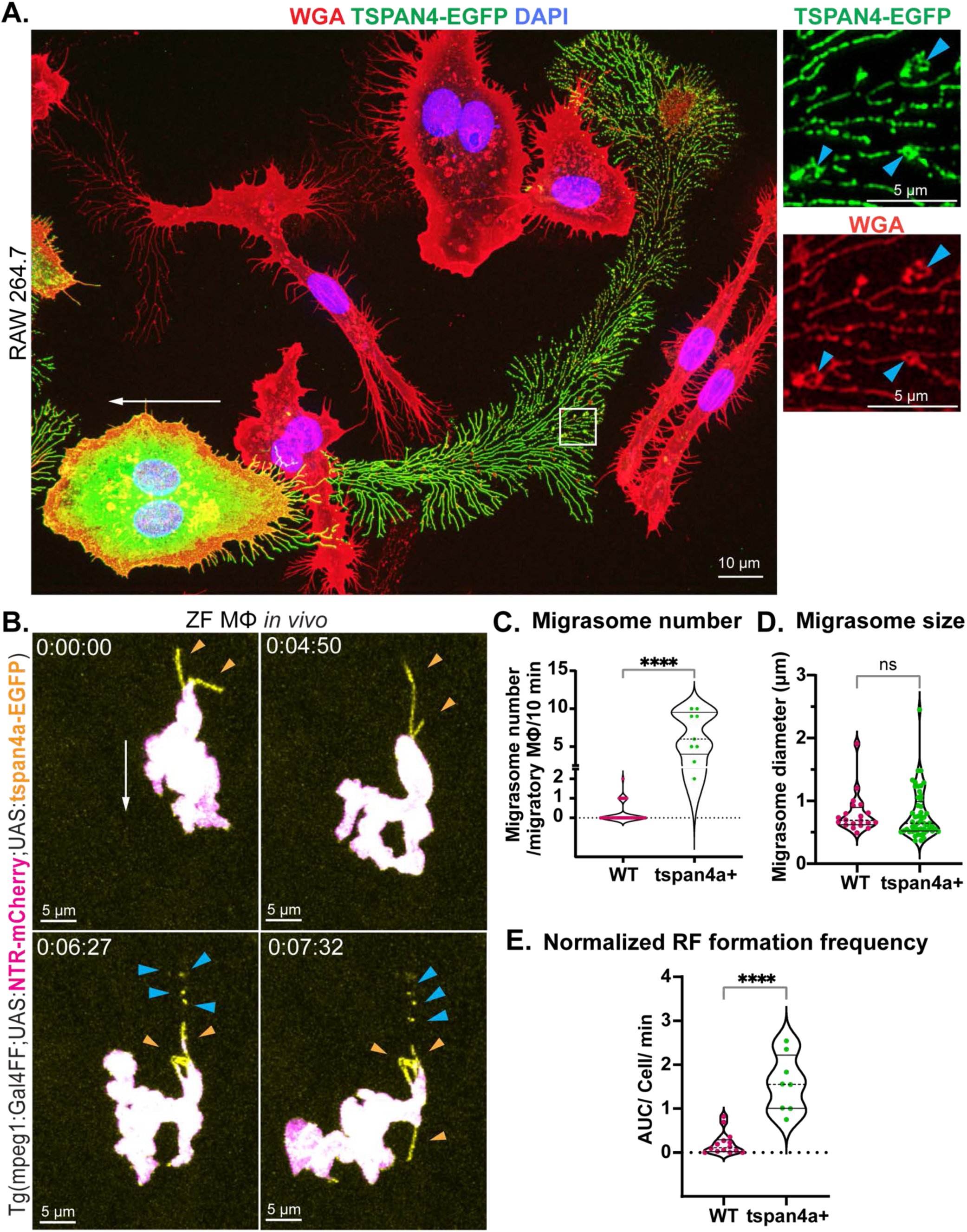
TSPAN4-EGFP marks macrophage retraction fibers and migrasomes and promotes their formation, *in vitro* and *in vivo*. **A.** TSPAN4-EGFP (green) transfected RAW macrophages, fixed after CSF-1 treatment and cells stained with Alexa Fluor 647-WGA (red) and DAPI (blue). Blue arrowheads indicate migrasomes; white arrow indicates cell migration direction. **B.** Timelapse series of a Tg(mpeg1:Gal4FF;UAS:NTR-mCherry;UAS:tspan4a-EGFP) macrophage *in vivo*, displayed as merged (white, MIP) from mCherry (magenta) / EGFP (yellow) channels. Note overlapping NTR-mCherry and tspan4a-EGFP expression in the macrophage soma (Mf), and tspan4a-EGFP-rich RFs (t=0:00, min:sec). These are flexible (t=4:50), bulge as tspan4a-EGFP-rich migrasomes form (t=6:27) and fragment (t=7:32) as the RF and migrasomes separate. The white arrow indicates direction of migration. Blue arrowheads and orange arrowheads indicate migrasomes and RFs respectively. See **Video 3**. **C.** Number of migrasomes formed by WT and tspan4a+ macrophages *in vivo* (normalized to number formed per 10 min). tspan4a+ macrophages form more migrasomes. n: WT=38, tspan4a+=9 cells. **D.** Size of migrasomes formed by WT and tspan4a+ macrophages *in vivo* are the same at this resolution. n: WT=20, tspan4a+=52 migrasomes. **E.** Parameter AUC/cell/min derived from RF number from per cell for tspan4a+ (n=8 cells) and WT (n=14 cells) macrophages (**S2A** and **S2B Fig**), an integrated parameter that is proportional to RF number, weights each cell equally and normalizes for different video lengths, demonstrating significantly increased RF formation by tspan4a+ macrophages. Statistics: **C** and **D**. Mann-Whitney test. **E**, Kolmogorov-Smirnov test. **** *p*<0.0001, ns=not significant. Data for **B-D** from MIP images, Zeiss LSM980 confocal microscope. **A** and **B**, scalebars as indicated.

Expression of the tspan4a-EGFP reporter in ZF4 cells enhances their formation of enlarged migrasomes (**S2D Fig**). We confirm that the cell autonomous overexpression of the TEM structural component TSPAN4/tspan4a in murine and zebrafish migrating macrophages promotes and enhances their formation of trails of RFs and migrasomes.

### Both migrasomes and RFs are sources of adherent and released extracellular vesicles

Live imaging of macrophages expressing TSPAN4-EGFP *in vitro* reveals the *de novo* formation of migrasomes along RFs. TSPAN4-EGFP is progressively concentrated upon packing into individual growing migrasomes, which increase in size and fluorescence intensity (**S3 Fig**), mirroring the process described in NRK cells (11). In migrating TSPAN4-EGFP-labeled RAW cells, live imaging shows migrasomes ranging in size, averaging 2.6 μm diam (**Fig 3A)** with retention times on RFs extending from 0.5 to 8 hrs, averaging 3.5 hrs (**Fig 3B**). Selected frames from a representative movie document the stable retention of a migrasome recorded on a RF for >6 hrs (**Fig 3C**). Other frequent fates are recorded here in movie frames, including, migrasomes detaching from RFs and dispersing after ∼1 hr (**Fig 3D**), new extracellular vesicles splitting off migrasomes (**Fig 3E**) and migrasomes being transferred from an RF into a neighboring cell (**Fig 3F**).

**Figure 3.**
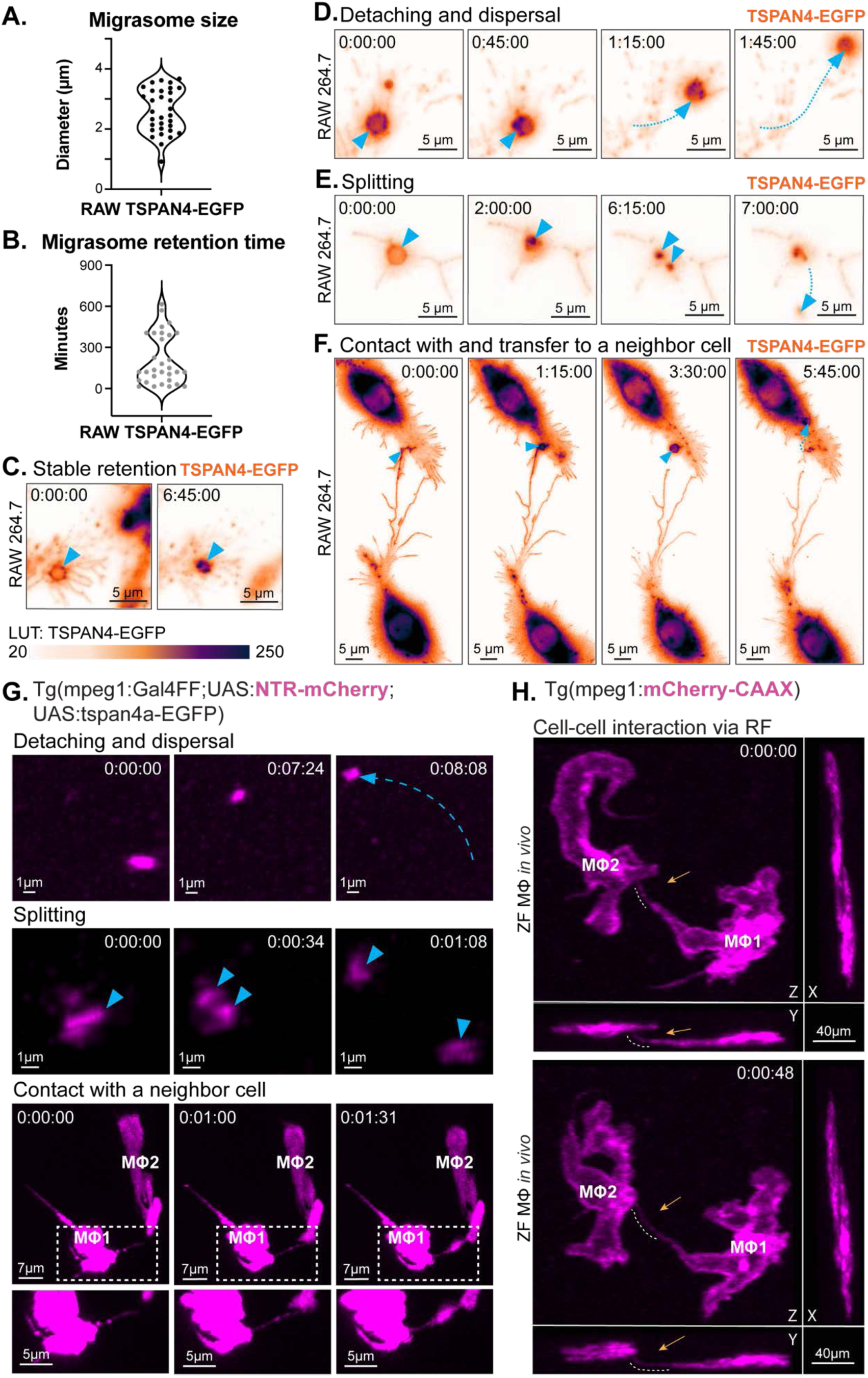
Formation and diverse fates of migrasomes *in vitro* and *in vivo*. **A.** Migrasome size distribution from TSPAN4-EGFP expressing RAW264.7 macrophages. Maximum diameter of maturing migrasomes measured manually in FIJI (n = 32 migrasomes). **B**. Migrasome retention time recorded and calculated using live imaging of TSPAN4-EGFP RAW264.7 cells in CSF-1,16 hrs. Migrasome fates demonstrated (n=31 migrasomes). **C**-**F**: Migrasome behaviors from TSPAN4-EGFP macrophages (CSF-1 for > 16 hrs, imaging interval = 15 min). LUT: inverted gem for TSPAN4-EGFP. Time stamp, hr:min:sec. **G.** Macrophage migrasome behaviors demonstrated *in vivo* in the Tg(mpeg1:Gal4FF;UAS:NTR-mCherry;UAS:tspan4a-EGFP) zebrafish reporter line. Blue dotted arrow in upper panel indicates direction of floating. Blue arrowheads in middle panel indicate the migrasome that has split in two. Lower panel insets highlight point of contact with a neighbor cell. **H**. Intercellular contact in the Tg(mpeg1:mCherry-CAAX) line. White dashed lines and orange arrows indicate the point of RF cell-cell contact. **G-H,** images from Zeiss LSM980 confocal. Only mCherry MIP channel shown. **A-H**, Scalebars as indicated.

Intravital imaging in the zebrafish tspan4a-EGFP-expressing line demonstrate the same migrasome behaviors *in vivo*. Macrophage migrasomes detaching from RFs (**Fig 3G top panel**) and migrasome splitting (**Fig 3G middle panel**) are shown. Macrophage-to-macrophage communication via RFs decorated with migrasomes is depicted in **Fig 3G lower panel**. Direct intercellular communication between macrophages is not an artifact of the tspan4a-overexpression reporter since they are also recorded in the mCherry-CAAX reporter line (**Fig 3H** shows two mCherry-CAAX macrophages tethered together by an RF extending between them). Thus, following varied periods of retention on RFs, macrophage migrasomes themselves can be released into the milieu or exchanged with other cells as extracellular vehicles.

We also examined the fates of RFs themselves in macrophages undergoing migration. Brightfield live imaging of RAW264.7 macrophages over 40 hrs reveals sub-populations of stationary cells and cells undergoing random migration in two distinctive patterns: ‘migratory cells’ moving along sustained directional trajectories and ‘oscillatory cells’ moving back and forth over the same short path (**S4A Fig** and **Video 4**). At higher resolution with confocal live imaging, migratory cells moving unidirectionally deposit RF and migrasome trails at their rear, while oscillatory cells moving back and forth switch their operational rear to always deposit trails behind them on their alternating trajectories (**Fig 4A** and **Video 5** and **6**). Thus, RF deposition is a polarized spatially-defined feature of migrating macrophages that is preserved across varied patterns of macrophage motion.

**Figure 4.**
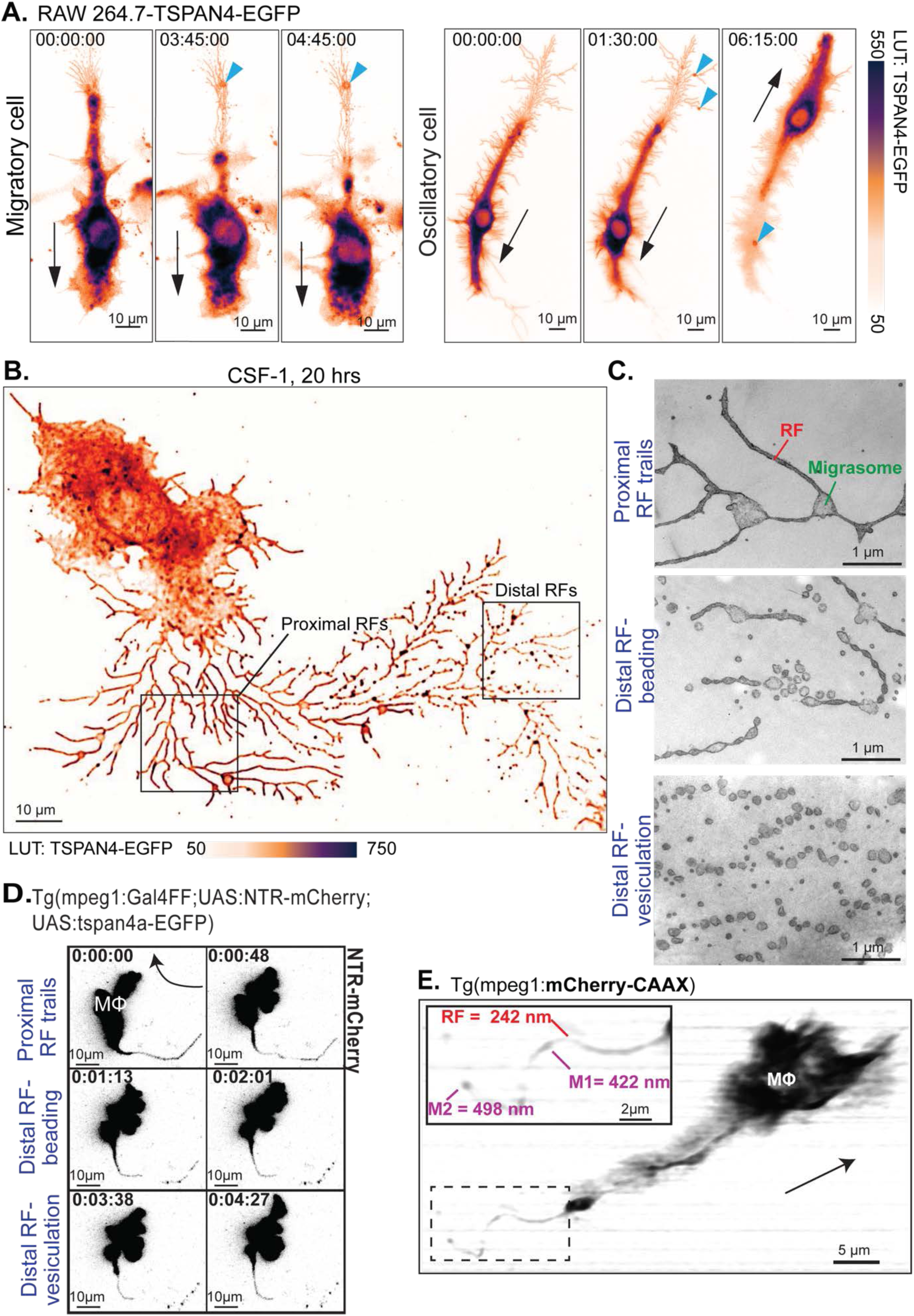
Maturation of macrophage RFs and migrasomes *in vitro* and *in vivo*. **A.** Fluorescence live imaging of TSPAN4-EGFP RAW264.7 cells on FN-coated substrates, recorded for 16 hrs at 15-min intervals on Zeiss Axio Observer 7. Migrasome formation (blue arrowheads) demonstrated in ‘migratory macrophage’ (left panels), and an ‘oscillatory macrophage’ (right panels). Black arrows indicate cell migration direction. See **Videos 5** and **6**. LUT: inverted gem for TSPAN4-EGFP. **B.** Single frame from live imaging of migrating RAW TSPAN4-EGFP cells captured on a 3i LLSM, proximal RFs and distal RFs (labeled boxes). LUT: inverted gem for TSPAN4-EGFP. **C.** RAW264.7 cells treated with CSF-1 for 16 hrs, fixed, resin embedded and thin sectioned for transmission EM to view RF and migrasome trails. **D.** RF and migrasome formation by a Tg(mpeg1:Gal4FF;UAS:NTR-mCherry;UAS:tspan4a-EGFP) zebrafish macrophage *in vivo*. Stills show RFs (t=0:00-0:48 min:sec) maturing to form beaded strings of migrasomes (t=1:13-2:21), which fragment to form dissociated migrasomes (t=3:38-4:24). Black arrow indicates cell migration direction. Zeiss LSM980 confocal MIP images, mCherry channel only. See **Video 7**. **E.** RF and migrasomes formed by a macrophage from the bioneutral Tg(mpeg1:mCherry-CAAX) reporter line, live imaged by LLSM, for precise measurement of native dimensions as labelled in inset. MIP images, Zeiss LLS7, mCherry channel only. Black arrow indicates migration direction. **A**-**E**, scalebars as indicated.

In addition to deploying migrasomes, macrophage RFs themselves are also a source of vesicles over time. Migrating TSPAN4-EGFP macrophages observed by LLSM show intact proximal RFs are contiguous with the cell body, but distal RFs, deposited earlier, start to fragment and detach from the moving cell (**Fig 4B**). By transmission electron microscopy (EM), the proximal RFs are confirmed as intact tubes (60-120 nm width, tapering from their base to distal extremity), with migrasome bases (∼520 nm diam) at branching junctions. The distal RFs disintegrate in phases, first by beading into strings of vesicles (**Fig 4C**), a process described as ‘pearling’ for the formation of extracellular vesicles from other membrane protrusions (25). Beaded RFs then break up further into individual vesicles (70-250 nm diam), which can be matrix-bound or float away (**Fig 4C, S4B Fig**). While Wang et al (18) have described a population of homogenous vesicles arising from RFs they named retractosomes, our results instead show highly varied vesicles and fragments appearing over time as a result of RF breakdown.

The distinctions between proximal and distal RFs are also evident in zebrafish macrophages *in vivo* **(Figs 4D and 4E).** Distal RF beading and vesiculation are recorded over time (**Fig 4D and Video 7**) in tspan4a-EGFP macrophages. Furthermore, in zebrafish macrophages marked by the biologically-neutral mCherry-CAAX reporter (**Fig 4E**), RF fragmentation is also seen, indicating that this dynamic sequence of macrophage RF maturation and vesiculation is an authentic physiological feature. Thus together, *in vitro* and *in vivo* both macrophage RFs and migrasomes provide initial arrays of matrix-adherent membrane trails but over time these membranes detach or are dispersed as extracellular vesicles.

### Recycling endosomes at the rear of migrating cells launch polarized trafficking into RFs

Establishing that the RF trails are always produced at the rear of migrating macrophages introduces a conundrum for how these motile cells regulate their intracellular membrane flow. Typically, in migrating cells exocytic membrane traffic is expected to flow forward to form the lamellipodia at the leading edge. This polarized traffic is coordinated by recycling endosomes (RE) positioned towards the front of the migrating cells which also serve to recycle leading edge integrins (26, 27). But, using LLSM in live cells, we record the active transport of cytoplasm and organelles into proximal RFs at the cell rear for the expansion of RFs (**S5A Fig, Video 8**). To examine REs, cells transfected with the recycling endosome marker Rab11a-mCherry show the perinuclear location of REs in stationary cells, but in migrating cells the Rab11-labeled REs are present in clusters at both the front and rear of the directionally-polarized cells (**Fig 5A**). Expression of another RE-associated GTPase, Rab8a shows a similar rear accumulation of RE membranes at the base of RFs and both Rabs11 and 8 can also be seen in migrasomes (**S5B and S5C Fig**).

**Figure 5.**
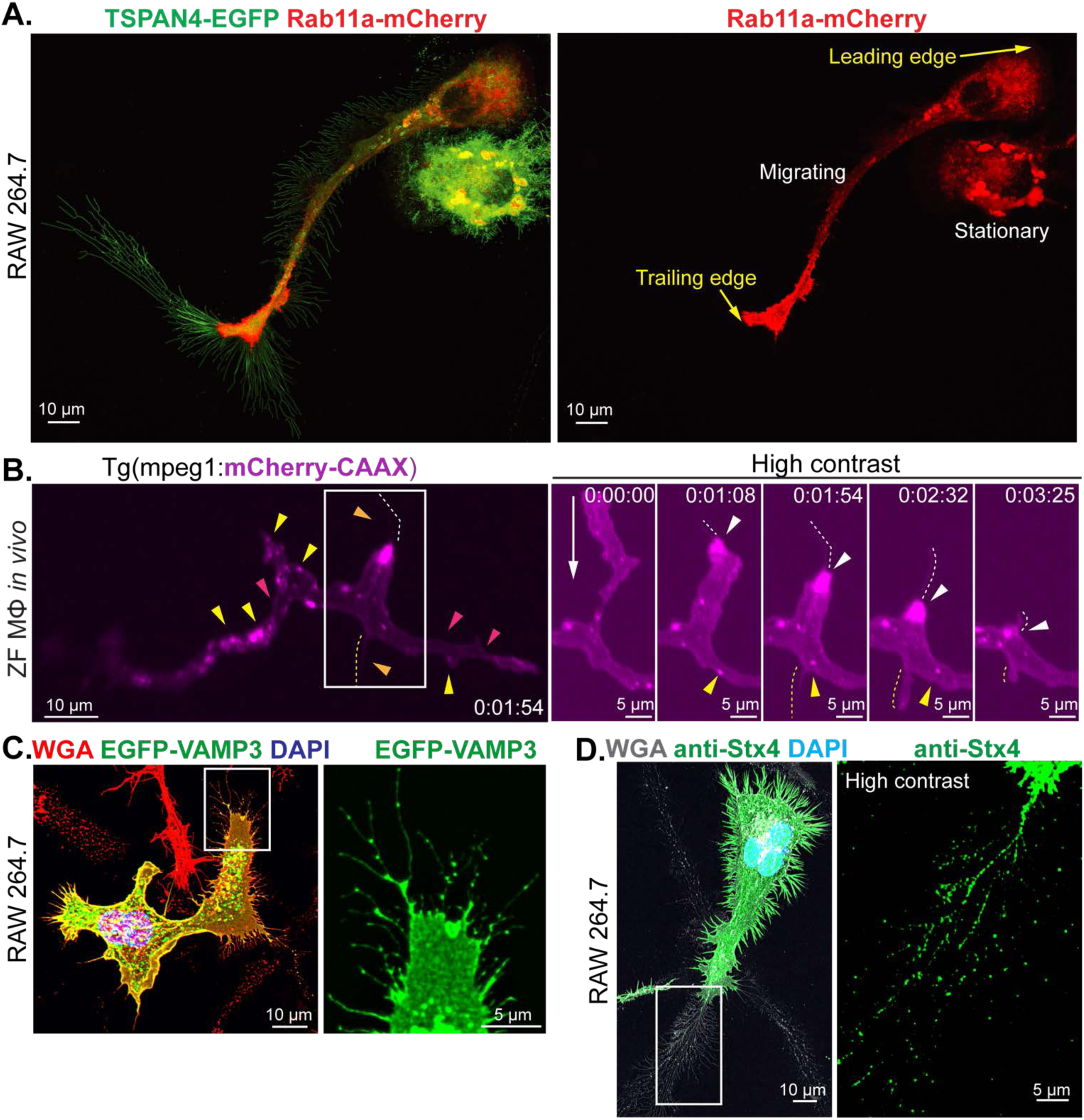
Localization of recycling endosomes and SNAREs in migrating macrophages *in vitro* and *in vivo*. **A.** TSPAN4-EGFP (green) and Rab11a-mCherry (red) cotransfected in RAW264.7 cells on FN-coated MatTek dishes for live imaging on Zeiss LSM880 airy scan confocal microscope. Trailing edge and leading edge of migrating macrophage indicated by yellow arrows, adjacent to a stationary cell. **B.** Recycling endosomes in zebrafish macrophage *in vivo*, targeted positioning near to the base of retraction fibers. Left panel is an overview of a Tg(mpeg1:mCherry-CAAX) macrophage at t=1:54 min:sec. Note the labeling of the outer cellular membrane, but also of multiple freely mobile intracellular vesicles throughout the cell (see **Video 9**), which delineates the external cytoplasmic membrane (pink arrowheads), internal membrane-lined vesicles (yellow arrowheads) and membrane-delineated RFs (dashed orange lines). The boxed area highlights one trailing arm that is followed in the time series of frames on the right. t=0:00 (min:sec), baseline appearance; t=1:08-2:32, progressive retraction of an extended protrusion towards the central macrophage soma, with condensation of the membrane reporter at its base (white arrowheads) as a RF forms and elongates (dashed yellow or white line and orange arrowheads; t=3:25, RF has disappeared and membrane signal at base is dispersing (white arrowheads). On the other side of the cell a smaller RF forms at t=1:54 (yellow dashed lines); as this RF forms and disappears, a small freely mobile vesicle (yellow arrowheads) moves to its base and then moves away. MIP image, Zeiss LLS7 LLSM. Long arrow indicates migration direction. **C.** EGFP-VAMP3 (green) transfected RAW264.7 cells in the presence of CSF-1, fixed and stained with Alexa Fluor 647-WGA (red) and DAPI (blue). **D.** RAW264.7 cells CSF-1 and LPS treated and fixed for Stx4 immunostaining (rabbit anti-Stx4 with a secondary antibody, Alexa Fluor 488-anti-Rabbit IgG, green) also stained with TMR-WGA (LUT: Grays) and DAPI (cyan). Imaged on upright wide field fluorescence microscope Zeiss AxioImager with Apotome2. **A-D**, scalebars as indicated.

The mCherry-CAAX reporter used to label zebrafish macrophages *in vivo* labels all lipid membranes of the cell and is enriched in REs. LLSM shows that in migrating zebrafish macrophages *in vivo*, multiple, freely-mobile mCherry-CAAX-marked endosomes aggregate and pause at the operational rear of the cells, adjacent to the bases of forming RFs (**Fig 5B, Video 1 and 9**). These labeled REs are always at the rear of these cells even as they change direction of movement, hence always defining the operational rear of these peripatetic cells.

To show that rear REs do support membrane traffic into the RFs, we examined RE-derived carrier vesicles, which in macrophages includes vesicles using the SNARE pair VAMP3-Syntaxin4 to move from REs to the plasma membrane (26, 28). Migrating macrophages expressing EGFP-VAMP3 label RFs and vesicles in RFs (**Fig 5C**) and Syntaxin4 (Stx4) immunostaining is found on the cell surface and along RFs (**Fig 5D**). Hence the necessary Rab and SNARE machinery is in place at the rear-facing pole of migrating macrophages for traffic to move directly or via REs into RFs and migrasomes and for formation of the RFs themselves. Since REs are known secretory hubs (29, 30), rear REs offer potential for exocytic trafficking into the RF trails.

### The transmembrane adaptor SCIMP is a functional TEM component of macrophage RFs and migrasomes

Transmembrane, palmitoylated adaptors of the pTRAP family also reside in TEMs (31). The pTRAP SCIMP is primarily expressed in macrophage lipid-rich membrane domains on filopodia, ruffles and in macropinosomes where it functions as a signaling adaptor for immune receptors (20, 32, 33). A significant finding here is that in migrating, LPS-activated macrophages, SCIMP is also seen on both RFs and migrasomes, revealed by immunostaining of endogenous SCIMP (**Fig 6A**). Transiently expressed Halo-SCIMP (**Fig 6B** and **S6A Fig**) colocalizes with TSPAN4-EGFP along RFs and on migrasomes. The association of SCIMP with RF trails is confirmed *in vivo* by imaging showing that a scimp-EGFP reporter targeted for expression in zebrafish macrophages localizes throughout macrophage soma and also on their associated migrasomes (**Fig 6C**). Thus, SCIMP is newly identified as a component of macrophage RFs and migrasomes.

**Figure 6.**
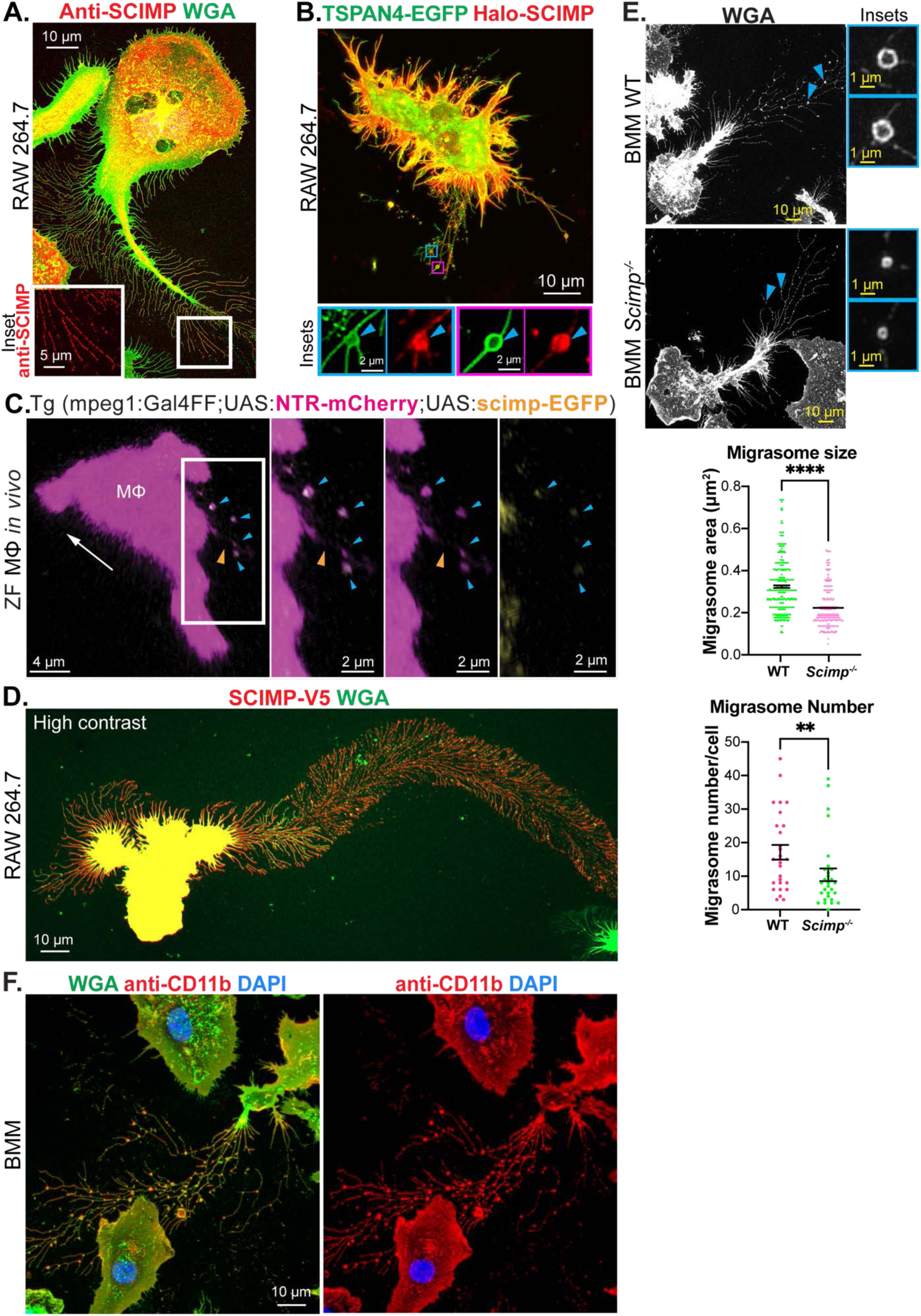
Tetraspanin-enriched microdomain molecules in macrophage RF trails *in vitro* and *in vivo*. **A.** RAW264.7 cells treated with CSF-1 and IFN-**γ** overnight and LPS for 4 hrs before fixation. Rabbit anti-SCIMP immunostaining visualized with a secondary antibody conjugated with Alexa Fluor 594 (red) and cell staining with Alexa Fluor 488-WGA (green) and DAPI (blue). High contrast inset shows SCIMP labeling along RFs. **B.** TSPAN4-EGFP (green) and Halo-SCIMP coexpressed in CSF-1 treated RAW264.7 cells colocalize in filopodia, RF, and migrasomes. Blue arrowheads in insets show migrasomes. **C.** scimp-EGFP reporter in zebrafish macrophages and macrophage migrasomes *in vivo*. Tg(mpeg1:Gal4FF;UAS:NTR-mCherry;UAS:scimp-EGFP) macrophage, displayed as merged (white) / mCherry (magenta) / EGFP (yellow). Orange arrowheads indicate RFs; blue arrowheads indicate migrasomes. White arrow indicates direction of migration, determined from timelapse video context. MIP image, Zeiss LSM980 confocal microscope. **D.** RAW264.7 cells expressing SCIMP-V5 fixed and immunostained with anti-V5 and Alexa Fluor 594 (red) secondary antibody, cell staining with Alexa Fluor 488-WGA (green). **E.** BMMs derived from wild-type (WT) and *Scimp^-/-^* mice treated with CSF-1 for 24 hrs and LPS, then fixed and stained with Alexa Fluor 488-WGA. Migrasome area and numbers per cell measured based on the WGA staining with FIJI ‘Measurement’ function. Representative migrasomes depicted by blue arrowheads and matching insets. n for migrasome area measurement: WT=457, *Scimp*^-/-^=286 migrasomes. n for migrasome number per cell: WT = 28, *Scimp*^-/-^ = 29 cells. Data is from 3 independent experiments and presented as mean ± SEM. Mann-Whitney test, **** *p* < 0.0001; ** *p* < 0.01. **F.** Immunostaining of CD11b in fixed, CSF-1 treated BMMs with a CD11b-specific antibody, an Alexa Fluor 594, (red) secondary antibody and cell staining with Alexa Fluor 488-WGA (green) and DAPI (blue). **A-F**, scalebars as indicated.

Similar to the effect of overexpressing TSPAN4-EGFP (**Fig 2A**), the overexpression of SCIMP-V5 produces extensive RF trails in migrating RAW264.7 macrophages (**Fig 6D**). We showed previously that BMMs from WT mice produce SCIMP but the SCIMP protein is absent in BMMs from *Scimp*^-/-^ mice (33). Here, in WT and *Scimp*^-/-^ BMMs, both RFs and migrasomes are produced by migrating cells (**Fig 6E**) and although the migrating *Scimp*^-/-^ BMMs have more variability in the amount of RFs formed, a consistent trend was not observed. However, the migrasomes produced by *Scimp*^-/-^ BMMs are significantly reduced in both number and size compared to those made by WT mouse BMMs (**Fig 6E)**. Thus, like TSPAN4, SCIMP represents a structural element of TEMs required for migrasome, and likely for RF production. Unlike TSPAN4, which is a more general component of these membranes, SCIMP is significant as a newly identified, inherent macrophage constituent of RFs and migrasomes in migrating cells.

Our screening revealed additional TEM components present in macrophage RFs and migrasomes. The distinctive macrophage surface integrin and inflammatory mediator CD11b is prominently stained in migrasomes and along RFs (**Fig 6F**). Other integrins such as the matrix-binding ⍺5 integrins can also be detected in the RF trails (**S6B Fig**). Filipin staining of cholesterol along RFs and RF fragments and enriched in migrasomes (**S6C Fig 6C**), is consistent with the cholesterol-rich nature of TEMs and migrasomes (8). Interestingly, EM images also depict outwardly-extruding double membrane buds (50-80 nm) emerging from the external face of the bilayer in RFs and migrasomes (**S6D Fig**). Protrusion-derived buds of similar size were previously described as vehicles for shedding excess membrane cholesterol from trailing membranes in macrophages (34). Migrasomes and RFs require further testing as possible sources for cholesterol shedding. Taken together, the presence of SCIMP and CD11b hint at roles for RFs and migrasomes in inflammatory responses, since these proteins have each been implicated in regulating cytokine outputs in activated macrophages (19, 21).

### Inflammatory cytokines synthesized by migrating macrophages are delivered to RFs and migrasomes

Activated macrophages traffic and secrete inflammatory cytokines in response to pathogenic and inflammatory stimuli. The pathways established for the RE-mediated trafficking and cell surface release of inflammatory cytokines such as interleukins (IL-6, IL-10) and tumor necrosis factor alpha (TNF), have relied to date on studies in stationary cells (28, 30, 35, 36). Based on the evidence presented above and in other studies (16, 17), RFs and migrasomes present another potential secretory site in migrating cells. In transfected, LPS and CSF-1 activated, migrating RAW cells, recombinant IL-6-EGFP and IL-10-EGFP can each be seen as perinuclear Golgi labeling and detected as faint staining carried in RFs and in migrasomes (**S7A and S7B Fig**). Thus, in addition to their release from the cell body, these two soluble cytokines can be trafficked into RFs and migrasomes in migrating cells.

The proinflammatory cytokine TNF is produced as a transmembrane precursor (pro-TNF) which is trafficked via the Golgi and REs to the cell surface, where cleavage by tumor necrosis factor alpha converting enzyme (TACE) releases the pro-TNF ectodomains as the soluble cytokine (37). In stationary macrophages, TNF is delivered to membrane ruffles, such as those around phagocytic cups (28). Upon examining migrating RAW cells transfected with EGFP-tagged TNF, the cytokine appeared along RFs and in migrasomes, as initial evidence that RF trails may represent a new migration-dependent site for TNF delivery (**S8A and S8B Fig**). In migrating RAW cells and BMMs, the LPS-induced *de novo* synthesis and trafficking of endogenous pro-TNF is further tracked by immunostaining (**Fig 7**). To retain and detect pro-TNF delivered to the cell surface, cells are incubated with the TACE inhibitor, TAPI (38). Immunostaining of pro-TNF is initially detected in the perinuclear Golgi complex by 30 min post addition of LPS (**Fig 7A** right panels), pro-TNF is then progressively delivered to cytoplasmic vesicles and the REs and it appears in RFs by 1-2 hrs (**Fig 7A**). There is a similar distribution of pro-TNF immunostaining in LPS-activated, migrating BMMs which have pro-TNF staining on the cell surface at the leading edge and in RF trails of migrating cells (**Fig 7B**). Pro-TNF is also detected in some migrasomes and along RFs (**Fig 7C**). In RAW cells, TNF staining is present on RFs and on migrasomes including those that have detached from the cell but remain adhered to the matrix (**S8C Fig**). Additionally, we show that TNF is trafficked into RFs via the rear REs, based on containing of endogenous TNF with tdTomato-Rab8a and coimmunostaining of TNF and Syntaxin4 in RFs (**S8D and S8E Fig**). Together, these results newly show that RFs and migrasomes in migrating cells are a key destination for the delivery of newly synthesized pro-TNF.

**Figure 7.**
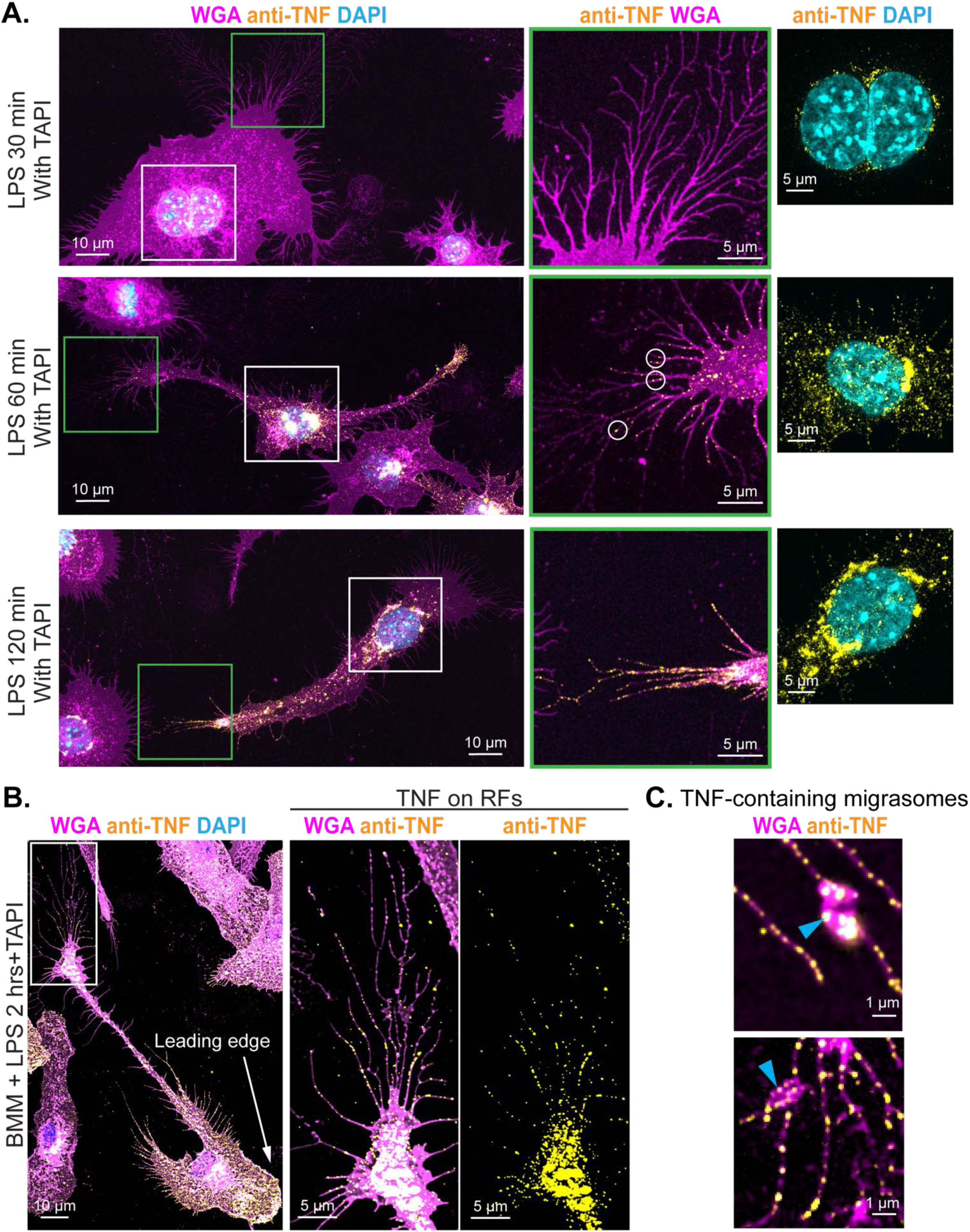
Newly synthesized tumor necrosis factor (TNF) delivered to macrophage RFs and migrasomes. **A-C.** Immunostaining of TNF (fluorescence of a secondary antibody Alexa Fluor 488 Donkey anti-rat IgG is shown, yellow LUT applied), cell staining with Alexa Fluor 647-WGA (LUT: Magenta) and DAPI (LUT: Cyan). **A.** RAW 264.7 cells treated with LPS (30, 60, or 120 mins) followed by TAPI incubation. Green-bordered insets show RFs studded with TNF staining (white circles). White-bordered insets show peri-nuclear staining of endogenous TNF. **B.** BMMs treated with LPS for 2 hrs, with TAPI for the last 30 mins. **C.** RAW264.7 migrasomes (blue arrowheads) with vesicular TNF staining. **A-C**, scalebars as indicated.

### RF trails from migrating cells serve to amplify local TNF presentation and secretion

While previous studies have focused only on migrasomes as vehicles for secretion (9, 17), here we reveal that both migrasomes and the RFs themselves are secretory sites for the presentation and release of TNF. Immunostaining without permeabilization of fixed cells detects abundant TNF on the surface of cell bodies and on the RFs and migrasomes in LPS-treated migrating macrophages (**Fig 8A**). Moreover, when TAPI is not added during incubations there is significantly less surface TNF along RFs and on migrasomes (**Fig 8B**), indicating that after temporary residence on these membranes, most pro-TNF is cleaved and released. Together these results newly demonstrate that the surfaces of RFs and migrasomes are sites for the delivery and release of TNF.

**Figure 8.**
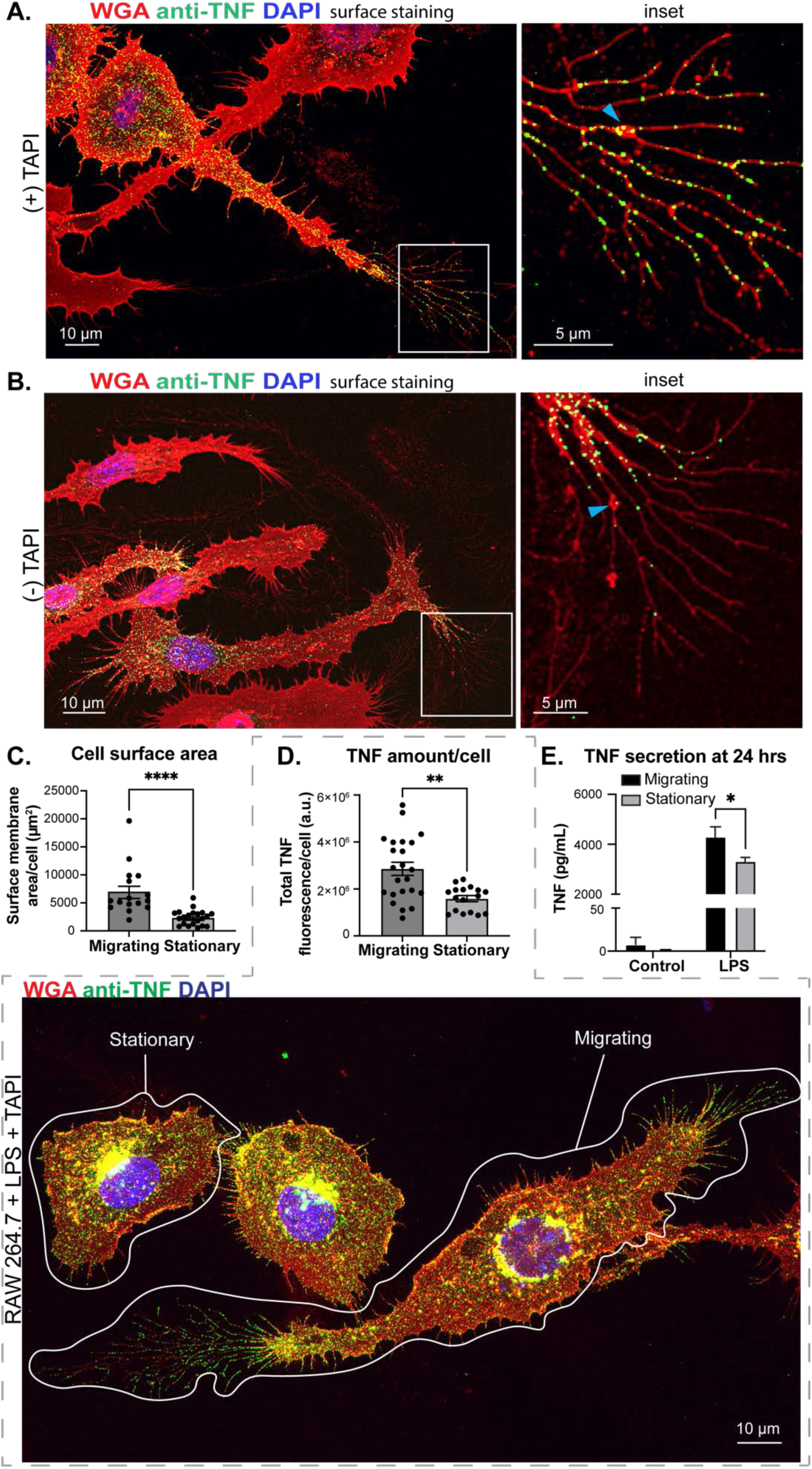
RF trails amplify TNF presentation and secretion in LPS-activated migrating macrophages. **A** and **B.** RAW264.7 cells treated with CSF-1 and LPS, with (+) or without (-) TAPI were fixed and immunostained without permeabilization to visualize cell surface TNF. TNF immunostaining with a secondary antibody (Alexa Fluor 488, green) and cell staining with Alexa Fluor 647-WGA (red) and DAPI (blue) stained cell membrane and nuclei respectively. Insets show endogenous TNF in RF trails. Blue arrowheads indicate migrasomes. **C.** Increased membrane surface area in migrating macrophages. 3D reconstruction of WGA-stained membranes in migrating cells (with RFs) and stationary cells (without RFs) yields total membrane surface area for each. Reconstruction and measurement were performed with Imaris. Pooled results of 4 independent experiments. Mann-Whitney test. n = 16 for migrating cells, n = 21 for stationary cells. **** *p* < 0.0001. **D.** Increased TNF presentation in migratory cells. RAW264.7 cells were activated with LPS for 2 hrs, with a 20-min delayed TAPI incubation. RAW264.7 cells were fixed, stained with Alexa Fluor 647-WGA (red) and DAPI (blue), immunostained against TNF with a secondary antibody conjugated with Alexa Fluor 488 (green). Image (bottom) shows segregation between stationary and migrating cells based on RF generation. Bar chart (top) shows the comparison of total TNF fluorescence intensity in migratory and stationary cells. Mann-Whitney test. Cell number n: migrating=23; stationary=17 across 3 independent experiments; ** *p* < 0.01. **E.** Increased TNF secretion by LPS-stimulated migratory macrophages. BMMs plated on tissue culture plates in CSF-1 and LPS (migrating) or on bacterial culture plates in LPS (stationary) as detailed in Material and Methods. Control groups were without LPS treatment. Data was from three independent experiments and presented as mean ± SEM. Two-way ANOVA. * *p* = 0.0214. **A**, **B** and **D**, Scalebars as indicated.

Quantitatively, the membrane surface area of RF trails potentially represents a significant expansion of sites for presentation and release of TNF in migrating cells. Measuring WGA-stained cells confirmed an increase in surface membrane in migrating cells (with RF trails) compared to stationary cells (no RF trails) (**Fig 8C**). Pro-TNF staining was compared in stationary and migrating cells within the same dishes (**Fig 8D**), revealing a concomitant increase in the amount of pro-TNF produced and displayed by migrating cells versus stationary cells in the same dishes (**Fig 8D**). Thus migrating cells contribute a significant pool of TNF that is initially presented in a spatiotemporal manner. The production and presentation of TNF is sustained over longer times, demonstrated in cells stained at 24 hrs post LPS (**S9A Fig**) and at this time point, migrating cells are secreting more soluble TNF into the medium compared to cultures of stationary cells (**Fig 8E**). Finally we observe that BMMs undergoing oscillating or short range migration over several days in culture, deposit carpets of overlapping RFs with migrasomes and staining of these dishes shows pro-TNF decorating these RFs in a concentrated manner (**S9B and S9C Fig**). Thus migrating cells have the capacity to be major, even dominant contributors, to the pool of spatiotemporally confined and dispersed TNF released by immune-activated macrophages.

### Migrating macrophages mediate TNF-dependent apoptosis in epithelial co-cultures

TNF deployment via RFs and migrasomes can be examined in more tissue-like environments using co-cultures of TSPAN4-EGFP macrophages and polarized IMCD3 kidney epithelial cells. The macrophages migrate through IMCD3 monolayers leaving RF trails in intercellular and subcellular spaces (**Fig 9A** and **B**). Treating cocultures with LPS induces TNF synthesis and the appearance of pro-TNF Golgi staining in the macrophages but not in IMCD3 cells under these conditions (**Fig 9C**). Upon addition of TAPI, pro-TNF immunostaining is visible for up to 24 hrs on the RFs deployed throughout the cell layer (**Fig 9C**).

**Figure 9.**
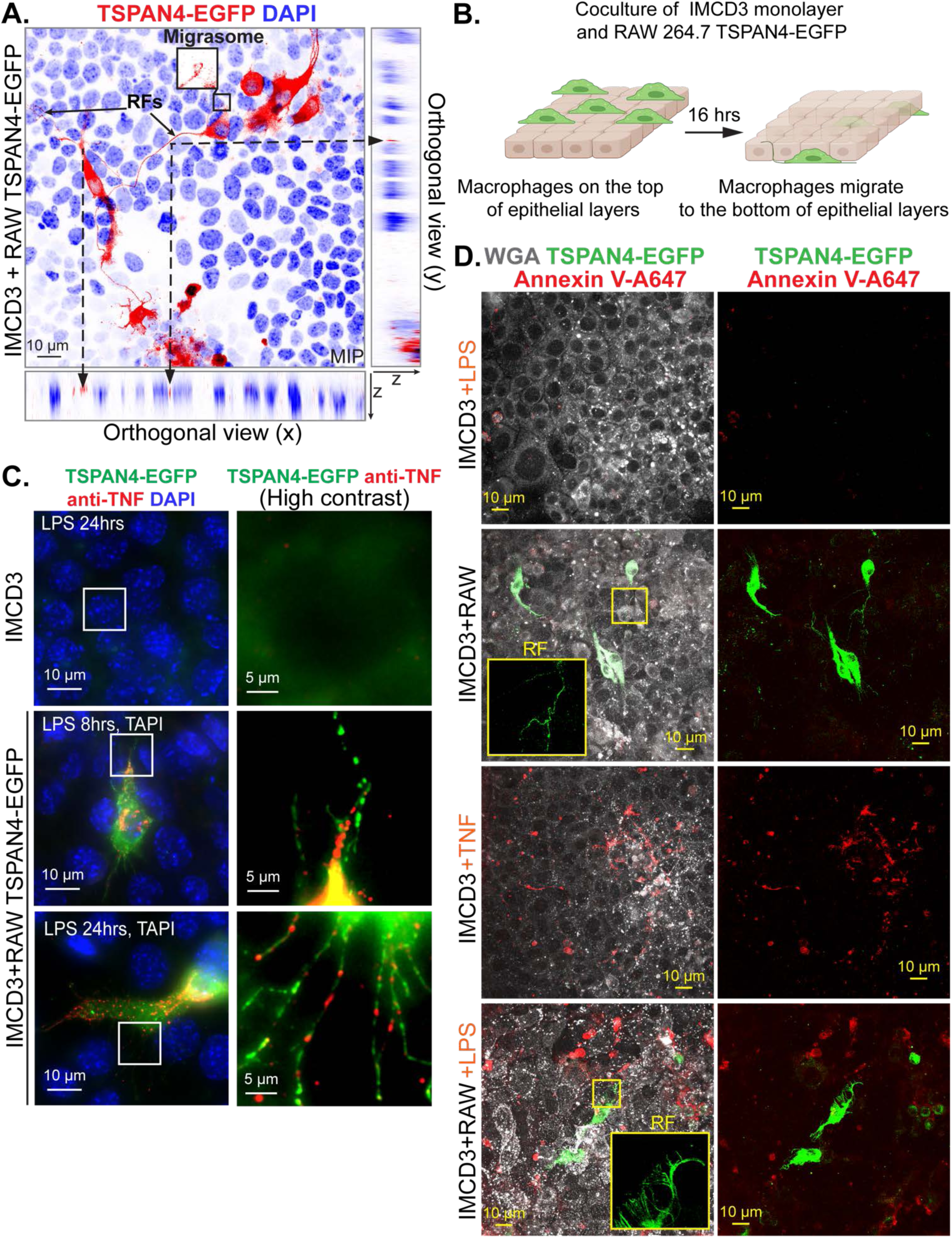
Migrating macrophages deployed TNF via RFs and mediate apoptosis in epithelial cocultures. **A-B.** Macrophages distribute throughout the epithelial monolayer. RAW264.7 TPSAN4-EGFP cells were replated on preformed IMCD3 monolayers and cocultured in RPMI full medium for 16 hrs with CSF-1. **A.** Cocultures fixed and stained with DAPI. Fluorescence imaging of z-stack presented as MIP with orthogonal views representing xz and yz planes. TSPAN4-EGFP positive macrophages (LUT: inverted Red) and all cells were stained with DAPI (LUT: inverted Blue). Corresponding points between MIP and orthogonal views connected by dashed arrows. RFs indicated with solid arrows. Migrasomes marked and zoomed in boxes with EGFP channel. **B.** Diagram of coculture of macrophages (RAW264.7 TSPAN4-EGFP, green) and epithelial monolayers (beige). Macrophages migrate from the top to the bottom of epithelial monolayers. **C.** Cocultures in LPS for 8 hrs or 24 hrs as indicated, last 30 mins with TAPI. Cocultures fixed, immunostained for TNF antibody (shown with an Alexa Fluor 647-conjugated secondary antibody, red), stained with DAPI (blue). Macrophages and their RFs were marked by TSPAN4-EGFP shown in green. Contrast enhanced insets present anti-TNF staining in RAW or IMCD3 cells (control). **D.** Cells were plated in MatTek dishes and treated with LPS (16 hrs), or TNF (8 hrs) as indicated, and imaged in the presence of Annexin V and WGA in culture conditions. Macrophages and their RFs marked by TSPAN4-EGFP in green. Alexa Fluor 350-WGA (LUT: Grays) staining shows IMCD3 monolayers. Apoptotic cells stained by Annexin V-Alexa Fluor 647 (red). **A**, **C** and **D**, Scalebars as indicated.

TNF binding to the receptor TNFR1 triggers downstream signaling in target cells with subsequent apoptosis providing a read-out for bioactive TNF secretion (39, 40). Direct addition of exogenous TNF to IMCD3 cell monocultures demonstrates the appearance of annexin V-stained apoptotic epithelial cells (**Fig 9D**). In other controls, co-cultures without LPS or the addition of LPS to monocultures of IMCD3 cells did not produce annexin V staining. In macrophage and IMCD3 cocultures treated with LPS, the release of TNF from migrating macrophages with RF trails, induces annexin staining of apoptotic epithelial cells (**Fig 9D**). Thus we demonstrate that the deployment of TNF by migrating macrophages liberating TNF-laden RF trails and migrasomes has a physiological consequence for neighboring cells.

## DISCUSSION

This study characterizes the trails of membranous RFs and migrasomes left by macrophages in their wake during migration *in vitro* and *in vivo*. The results we obtain in fish embryos and in mouse cells, complement and extend features of RFs and migrasomes reported in the literature, revealing specific new features with particular relevance to macrophages, including identifying the pTRAP SCIMP as a structural component of RFs and migrasomes and identifying both migrasomes and RFs as sites for presentation and secretion of inflammatory cytokines. Whilst immune cells participating in anti-infective or inflammatory responses are often migratory, the contributions of migrating cells to inflammatory responses have until now been overlooked. Our findings establish that migrating macrophages are active proponents of inflammatory responses, moreover they are customized to contribute uniquely and significantly by leveraging their trails of RFs and migrasomes to enhance the retention and release of cytokines in a spatiotemporal fashion, for impact at local sites of inflammation.

We establish the *in vivo* equivalency of macrophage RFs and migrasomes analyzed in zebrafish and those more readily seen *in vitro* in cultured murine macrophages. The live imaging of migrating zebrafish macrophages, particularly using the biologically neutral mCherry-CAAX and NTR-mCherry reporters alongside the tspan4a-EGFP reporter, provides corroborating *in vivo* insights into the normal morphology and physiology of RFs and migrasomes, including the tspan4 (TSPAN4)-induced migrasome expansion and the detection of SCIMP as a new marker and structural component. Around a third of macrophages *in vivo* were seen to be making long RFs and while employing LLSM allowed us to capture their dynamics, we believe this imaging underrepresents the extent of the RF trails produced by migrating macrophages *in vivo,* warranting future deep tissue imaging approaches. We also employed a tissue-like setting using macrophage and epithelial cell cocultures *in vitro* (**Fig 9**), identifying macrophage RF deployment and TNF release among epithelial cells and extending this to show consequential TNF-induced epithelial apoptosis. Indeed it will be fascinating to resolve RF trails emerging from multiple immune cell types *in vivo* in tissues and at wound sites and our identification of macrophage markers, such as CD11b and SCIMP, is an important step towards this goal. The identification of RFs and migrasomes produced by migrating macrophages now adds to an inventory of novel and specialized morphologies detected using LLSM in migrating fish macrophages and neutrophils (41), highlighting the distinct physiology and functions of migrating versus stationary immune cells.

Overexpressing TSPAN4 in mouse macrophages or tspan4a in fish or ZF cells enhances RFs and migrasomes, specifically quantified as increases in RF frequency, migrasome number and size across these models (**Fig 2**). These findings are consistent with the elucidation (10, 11) that excess TSPAN4 increases the packing of TEMs into tetraspanin-enriched macrodomains (TEMAs) and increasing migrasome abundance, in contrast to tspan4a knockout in fish which reduces migrasome numbers (14). Here we reveal a new TEM component of RFs and migrasomes, the transmembrane pTRAP, SCIMP. Similar to TSPAN4, higher expression of SCIMP equates with more dramatic RF trails, while migrasomes produced by SCIMP knockout BMMs are reduced in size and number, suggesting that SCIMP contributes structurally to RF and migrasome TEMA formation to expand these structures. The increased expression of endogenous SCIMP (42) in immune activated and CSF-1 treated macrophages is consistent with a role for SCIMP in RF and migrasome formation in migrating cells. Interestingly, a possible mechanism for SCIMP’s expansion of TEMAs in RF trails is suggested by the behavior of a related pTRAP family member, LAT, which aggregates in lipid rafts in T cell membranes (43) where it has been imputed with a role in cholesterol packing in rafts to stabilize receptor signaling (44). It will be of interest to determine whether SCIMP could play a role in cholesterol loading for expansion of TEMAs in migrasomes and RFs, which are particularly enriched in sphingolipids and cholesterol (12) and have the appearance of shedding lipids in our EM studies (**S6 Fig**). Whether SCIMP also supports immune receptor signaling in the TEMAs of RFs and migrasomes in migrating macrophages is currently being investigated. Nevertheless, SCIMP now appears to have at least two proinflammatory roles which could contribute to defense and disease, firstly as a signaling adaptor for boosting the production of proinflammatory cytokines (21) and now secondly through formation of RF trails in migrating macrophages as platforms for enhancing inflammatory cytokine distribution. Finally, a recent study proposes a role for a secreted exosomal form of SCIMP as a chemoattractant in *E.coli*-but not LPS-activated macrophages, hence it remains to be seen in a broader context and if it is applicable across species for SCIMP’s variable extracellular domain(45).

The intracellular route for trafficking and secretion of newly-synthesized TNF in non-migrating macrophages has been well documented by us and others, wherein pro-TNF is delivered, via REs, to surface ruffles and filopodia for retention or TACE-mediated release (23, 28, 38). Other reports show that TNF can also be released from non-migrating macrophages in microvesicles (46). Now we find that during migration, macrophages use additional sites for the release of soluble cytokines and for the presentation and release of TNF. Migrating macrophages unexpectedly position Rab11a and Rab8a-associated REs at the rear of the cell as a hub for delivering membrane and cargo in RE-derived carriers into proximal RFs and migrasomes. The cognate SNAREs VAMP3 and Syntaxin4 are positioned for surface delivery of TNF in RFs and migrasomes, just as they are for delivery of TNF to lipid rich protrusions on the cell body (28, 38). Deploying TNF and other cytokines via spatiotemporal RF trails, fundamentally changes the current precept that soluble cytokines are released in a free-floating manner for unregulated systemic diffusion.

Migrasomes have previously been portrayed as highly uniform structures and as a well-defined, biochemical fraction (9, 14, 15), but our results emphasize that macrophage migrasomes are more inherently variable in size and appearance. Additionally, whereas RFs have been shown to form vesicles termed retractosomes (18), these have been overlooked as secretory sites (9). Our labeling shows that RFs themselves are prominent sites for direct TNF surface release and for vesiculation. Our live cell, EM and intravital images show macrophage RFs beading over time to form variable-sized vesicles and fragments, especially *in vivo*, where this process can be physically accelerated. Two key insights from our findings show how RFs and migrasomes are optimized for functions like cytokine release in macrophages, i) by deploying diverse arrays of extracellular vesicles whose inherent variation likely aids their interaction with multiple tissue elements and other cell types and ii) the dynamic and temporal nature of RFs and migrasomes as adherent platforms allows the local cytokine milieu to switch upon critical macrophage polarization from M1 to M2 outputs (47).Thus we propose that migrating macrophages generate variable and dynamic extracellular trails ensuring the greatest physiological versatility, temporal flavor and impact for signaling to other cells in multicellular and inflammatory environments.

Remarkably, quantification revealed that the trails of RFs and migrasomes provide migrating cells with enhanced ability to release cytokines compared to stationary cells. So, migrating cells may be the most active immune respondents, at least in terms of cytokine production, and certainly it will now be important to examine other, pro-and anti-inflammatory macrophage responses in the context of migrating cells. Due to the species-specific nature of TNF in fish, reagents are not available to confirm the role of macrophage RFs and migrasomes *in vivo* at this stage. However, previous studies have reported migrasomes as secretion sites for growth factors and other mediators in fish, chicken embryo and mice (14–16) and fibroblasts (17). In addition to being enhanced, the salient feature of cytokine release via RF and migrasome trails is the spatiotemporal nature of this dispersal. Having erstwhile soluble mediators corralled, either temporarily or over longer times, in tissue locales has the potential to locally amplify inflammatory and anti-infective responses, including by activating and recruiting other immune cells in the region. This may, for instance, be a key mechanism for locally amplifying inflammatory responses at sites of infection or injury, but more insidiously, it could also contribute to triggering or exacerbating acute cytokine storm or chronic inflammation. RFs are envisaged, and indeed seen *in vivo* in zebrafish macrophages, as forming long trails behind cells undergoing long-range migration, but findings support an additional scenario wherein the short range or oscillating migration of macrophages serves to deposit an interwoven carpet of shorter RFs (**S9B** and **S9C Fig**) which would act to further concentrate cytokine presentation and release at wound sites. Even after their release from RFs, migrasomes bearing TNF can be seen reattached to the surrounding matrix (**S8C and S9C Fig**) and TNF itself can reportedly bind to extracellular matrix molecules such as fibronectin (48). Thus, the deployment of adherent RFs and migrasomes provides macrophages with modular platforms for generating spatial cytokine gradients.

Taken together, the findings reported here provide new information about the deposition of RFs and migrasomes by migrating cells with a cell type (macrophage) specific focus *in vitro* and *in vivo*. These results reveal that immune cells simultaneously manage migration and cytokine secretion as part of the primary immune response in actual situations. The findings presented here fundamentally change current perceptions of inflammatory cytokine release in tissue environments where migrating, not stationary, macrophages are the most productive source of inflammatory cytokines and the cytokines themselves are not dispersed but instead displayed and released in a localized spatio-temporal fashion by retraction fibers and migrasomes. This work lays a foundation for further investigation of these migration-specific structures and immune functions in anti-microbial and inflammatory responses and more broadly in macrophage roles in development and disease.

## MATERIALS AND METHODS

### Murine primary cells and cell lines

Male and female mice of 8-12 weeks of age were housed with rodent chow and water *ad libitum* in groups of up to five animals per cage in a specific pathogen-free (SPF) facility with a 12-hr light/dark cycle. Mouse ethics and experimental protocols were approved by the University of Queensland (IMB/351/19). C57BL/6N mice from an in-house breeding colony were used as wild-type (WT) primary macrophages. *Scimp^−/−^* mice (strain C57BL/6N-Scimp^tm1a(KOMP)Wtsi^/Mmucd, RRID: MMRRC_049566-UCD) were described previously (33). Mouse bone marrow-derived macrophages (BMMs) were isolated as previously described (33). Briefly, primary mouse bone marrow cells harvested from femurs of C57BL/6 mice, were cultured in RPMI medium with 10% FBS, 2 mM L-glutamine, penicillin (20 units/mL) and streptomycin (100 ng/mL) and differentiated to bone marrow-derived macrophages (BMMs) by incubating in 75-100 ng/mL macrophage colony-stimulating factor (M-CSF, also referred to as CSF-1, ImmunoTools GmbH: 11343118) for 7-10 days and then used immediately for experiments. SCIMP expression in WT and its absence from *Scimp^-/-^* BMMs was established previously (33).

The RAW264.7 macrophage cell line obtained originally from ATCC (SC-6003) was passaged 2-3 times weekly and maintained in RPMI with 10% fetal bovine serum (FBS, Gibco: 10100147) and 2 mg/mL L-glutamine in 95% air, 5% CO_2_, 37°C under mycoplasma-free conditions. For stable expression of TSPAN4-EGFP, RAW264.7 cells were transfected with pEF6-TSPAN4-EGFP, put under selection in blasticidin (ThermoFisher Scientific: A1113903) at 2 μg/mL to generate stably expressing clonal cell lines, then removed from blasticidin the day before experiments.

To encourage cell migration and retraction fiber formation, cells were typically plated on glass coverslips pre-coated with fibronectin (FN, Sigma-Aldrich: F0895, 20 μg/mL), placed in 24-well plates and treated with CSF-1 (100 ng/mL, 6 hrs to overnight). To activate macrophages for cytokine production cells were treated with LPS (100 ng/mL) for the times indicated. To retain cell surface pro-TNF for immunostaining, cells were incubated with the TACE inhibitor TAPI (42), which was typically added 15 mins after LPS.

For co-culture experiments, IMCD3 (murine inner medullary collecting duct: mIMCD-3 line originally from ATTC) epithelial cells were plated first in MatTek dishes for cell death assays and live imaging or on glass coverslips in 24-well plates for immunostaining, maintained in complete DMEM, supplemented with 10% heat-inactivated FBS (v/v) and 2 mM L-glutamine at 37°C in a humidified incubator with 5% CO_2_. Confluent IMCD3 cells form epithelial monolayers with cell-cell junctions. TSPAN4-GFP expressing RAW264.7 cells were plated on epithelial monolayers at 0.2 × 10^6^ cells/mL with or without CSF-1. The coculture was maintained in RPMI full medium. Treatment and experiments were performed 16 hrs after RAW264.7 seeding.

### DNA constructs for murine protein expression *in vitro*

The sequence of mouse TSPAN4 cDNA was obtained from NCBI CCDS database. A gene block fragment containing the TSPAN4 sequence and restriction sites of EcoRI and BamHI was purchased from IDT Technologies (Singapore). The gene fragment was cloned into pEF6-EGFP-N1 vectors (adapted from pEF6/V5-His TOPO, ThermoFisher Scientific: K961020) through EcoRI/BamHI restriction sites as pEF6-EGFP-N1-TSPAN4 constructs. The mouse SCIMP full-length construct was amplified by PCR from cDNA and cloned into the pEF6/V5-His TOPO TA expression vector to yield a pEF6/V5-His-TOPO TA-SCIMP construct, previously described (21), used for SCIMP-V5 transfection. The construct was further modified by adding a gBlocks gene fragment containing the transmembrane signal peptide (TSP) and 2xMyc-tag and a HaloTag to the N-terminus of the SCIMP protein, generating a pEF6-TSP-Halo-SCIMP-V5 construct, previously published by (33). Additional constructs made in the Stow lab include pEF6-EGFP-VAMP3 (26), tdTomato-Rab8a (49), pEGFP-Rab8a (50), pEGFP-N1-IL-6 (51), pEGFP-C1-TNF (51), and pEGFP-N1-IL10 (35). Rab11a gene fragment was excised from a pEF6-EGFP-Rab11a construct (29) and cloned into pmCherry-C1 vector (Clontech: 632524) using BspEI and EcoRI restriction sites. TNF was also cloned into a pEGFP-C1 vector to produce an N-terminal GFP-tagged protein (Clontech) (51).

### Transfection of murine cells

For transfection with lipofectamine, RAW264.7 cells were seeded onto FN-coated coverslips in 24-well plates and MatTek dishes at a density of 0.05 × 10^6^ cells/mL. Following overnight recovery, Lipofectamine 2000™ (Thermo Fisher Scientific: 11668027, Australia) was used to transfect the cells according to the manufacturer’s instructions. Briefly, 1-2 μg DNA was mixed with Lipofectamine in Opti-MEM (Thermo Fisher Scientific: 51985034, Australia) and kept at room temperature for 5 min to form the DNA-Lipofectamine complex which was then added to cultures (125 μL per coverslip; 250 μL per MatTek dish). Cells were incubated with the complex for 2-4 hrs, refreshed with complete culture medium and maintained until use.

For electroporation, RAW264.7 cells were split and seeded at a density of 1 × 10^6^ cells/mL in a 10 cm Johns petri dish (Thermo Fisher Scientific). At confluence, cells were harvested and washed twice with serum-free medium. The cell suspension aliquoted into Gene Pulser/MicroPulser Electroporation cuvettes (Bio-Rad, 0.4 cm), each containing 2 × 10^7^ cells mixed with 10 μg (30 μL) of linear DNA. Electroporation was performed at 240 V and 1000 μF capacitance, cells were then washed with complete RPMI medium to remove debris and seeded onto tissue culture-treated dishes or coverslips. Blasticidin was added to the medium for selection and maintenance of stable cell lines, and was removed from the culture medium one day prior to experiments.

### ELISA assays

BMMs were replated on low-adhesion bacterial dishes to inhibit migration or on tissue culture plates supplemented with CSF-1 to promote migration on the 7th day of differentiation. Cells were then incubated with LPS (100 ng/mL) for 24 hrs under standard culture conditions. The medium was collected for ELISA to quantify TNF levels. Cells from both plate types were fixed and stained with WGA to assess the formation of RFs, which were observed only in migrating macrophages. The BD Biosciences Mouse TNF ELISA Set II (558534) and BD OptEIA™ TMB Substrate Reagent Set (555214), were used according to the manufacturer’s instructions. Pre-centrifuged medium collected from RAW cells incubated with or without LPS for up to 24 hrs was diluted in serum-free medium, incubated for 2 or more hrs in 96-well plates (Thermo Fisher Scientific: 439454) precoated with the capture antibody solution (1:250 in 0.1M sodium carbonate solution, pH 9.5), then blocked with assay diluent (10% FBS in PBS). Detection of bound cytokines was by sequential incubation with detection antibodies and Streptavidin-HRP conjugated reporter antibodies (1:250 in assay diluent each). Plates were washed 3-7 times between steps with a 0.05% Tween-20/PBS solution. Lastly, the reagent solution of tetramethylbenzidine (TMB) and hydrogen peroxide was added for a maximum of 30 min, and reaction was stopped using a 2N sulphuric acid solution. Readings were obtained using a Tecan Infinite M Plex plate reader at 450 nm with an absorption correction at 570 nm, and results processed and presented using GraphPad Prism (v10.0.2). The standard curve used a 4-parameter sigmoidal logistic curve.

### Fluorescence labeling and immunostaining of murine cells

Cultured cells fixed in paraformaldehyde (PFA, 4% in PBS) for 0.5-1 hr, were washed in PBS and then permeabilized (if required) with 0.1% TritonX-100 for 5 min (Sigma Aldrich), blocked with 0.5% bovine serum albumin (BSA) in PBS (Sigma Aldrich) and incubated sequentially in primary antibodies and fluorescently tagged secondary antibodies for 1 hr each with washing in between, or cells were incubated with fluorescent dyes while cocultures were incubated with primary antibodies for 2 days at 4°C and secondary antibodies together with fluorescent dyes for 3 hrs at RT. After final rinsing, all coverslips were mounted on Superfrost glass slides with ProLong Glass Antifade (Thermo Fisher Scientific) and sealed with nail varnish. Primary antibodies are monoclonal rat anti-TNF (BD Biosciences: 554416), rabbit Syntaxin4 (38), monoclonal rat anti-CD11b (Abcam: ab8878), anti-CD49e (integrin ⍺5, BD Biosciences: 55565), anti-V5 (Bio-Rad Laboratories Pty Ltd: MCA1360G), and rabbit anti-SCIMP (produced by the Walter and Eliza Hall Institute antibody facility, previously published (21)). Secondary antibodies are Alexa Fluor 488 Donkey Anti-Rat IgG (Invitrogen: A21208), Alexa Fluor 594 Goat anti-Rat IgG (H+L) Cross-Adsorbed Secondary Antibody (Invitrogen: A11007), Alexa Fluor 594 goat anti-rabbit IgG (H+L) Cross-Adsorbed (Invitrogen: A11012), Alexa Fluor 488 Goat Anti-Rabbit IgG (H+L) Cross-Adsorbed Secondary Antibody (Invitrogen: A11008), Alexa Fluor 594 Donkey Anti-Mouse IgG (H+L) Highly Cross-Adsorbed Secondary Antibody (Invitrogen: A21203).The blocking step was skipped for staining with membrane and nuclear dyes, including Wheat Germ Agglutinin (WGA) conjugated with Alexa Fluor 488 (Invitrogen: W11261), Tetramethylrhodamine (TMR) (Invitrogen: W849), or Alexa Fluor 647 (Invitrogen: W32466) as indicated and 4’,6-Diamidino-2-phenylindole dihydrochloride (DAPI, Sigma Aldrich: 32670). To stain cells with filipin (Sigma Aldrich: F-9765), fixed cells were incubated with 1.5 mg/mL of glycine in PBS for 10 min and then with 0.05 mg/mL filipin solution (diluted in 10% FBS/PBS) for 2 hrs. The coverslips were then washed with PBS and viewed under the microscope at 350 nm excitation. To visualize Halo-tagged recombinant proteins, cells expressing Halo-SCIMP-V5 were incubated with JF549 HaloTag^®^ ligand (kindly provided by Luke Lavis, Janelia Research Campus, VA) prior to imaging for 20 min at 10 nM. Live imaging was performed in the presence of HaloTag^®^ ligand. For recycling endosome labeling, cells were serum-starved for 30 min to deplete surface-bound transferrin and then incubated with Alexa Fluor 647 conjugated transferrin (25 μg/mL in full medium; Invitrogen: T23366) at 37°C for 12-15 min, followed by a 20 min chase before fixation. Fixed murine cells were imaged on a Zeiss AxioImager M2 upright microscope with Apotome2 and Axiocam 506 monochrome and color cameras using a Plan Apochromatic 63x NA 1.4 oil immersion objective; images were captured and deconvolved using Zeiss Zen Blue software.

### Apoptosis assay for murine cell cocultures

Apoptosis was detected with Annexin V-Alexa Fluor 647 staining in the cocultures of RAW264.7 and IMCD3 cells. Staining and imaging was performed as previously described (52). An annexin V binding solution consisted of HBSS (0.14 M NaCl, 0.005 M KCl, 0.001 M CaCl_2_, 0.0004 M MgSO_4_.7H_2_O, 0.0005 M MgCl_2_.6H_2_0, 0.0003M NaHPO_4_.2H_2_O, 0.0004 M KH_2_PO_4_, 0.006 M D-glucose, 0.004 M NaHCO_3_) containing an additional 0.005 M CaCl_2_ and 5% FBS. Cells were imaged in the binding solution with WGA-Alexa Fluor 350 at 10 μg/mL in culture conditions.

### Live imaging of murine cells *in vitro*

For live imaging, cells were plated on MatTek dishes and treated with or without CSF-1 at 100 ng/mL. Cells were cultured in Leibovitz’s L-15 medium (Invitrogen: 11415064) with 10% FBS and 2 mM L-glutamine at 37°C with 5% CO_2_ during imaging. Cells were imaged using aZeiss Axio Observer 7 inverted microscope with Axiocam 705 camera, or an LSM880 inverted confocal microscope with Airyscan both with incubation running either Zen Blue or Black software respectively. Live imaging was also carried out on a Lattice Light Sheet microscope (LLSM) (3i, Intelligent Imaging Innovations Inc., Denver, CO, USA). EGFP-expressing cells were excited with a 488nm laser using the following multi-bessel beam settings (44 beams, spaced 0.932mm apart with a cropping factor of 0.175), using the 0.493/0.55 annular mask. 3D volumes were acquired with a Z-step size of 0.495nm (after deskew 0.268 nm) using a Hammamatsu Orca Flash 4.0 sCMOS camera running Slidebook 6 software. Co-cultures were imaged using an inverted confocal microscope Zeiss LSM 880 with Airyscan with Plan Apochromat 40X Oil immersion lens using Zen Black software. For long term live imaging of cells grown on 24-well plates an Olympus Provi (Evident: CM30 kindly made available by Evident Australia) was used to record cell migration at 4X magnification with cells maintained within a CO_2_ incubator.

Image analysis was carried out on ImageJ2 / Fiji (53). LLSM live imaging was deskewed and deconvoluted for each time point in parallel on a high performance compute cluster running the Image Processing Portal. Live imaging data was viewed and cropped using Imaris, and deconvolved using Microvolution (Microvolution, LLC) plugin in Fiji. Time sequential maximum z-project was presented for all the live imaging. Images presented are also re-colored and/or inverted using Fiji where appropriate since there is no alteration of information in this process. Cell migration tracking on brightfield live imaging captured on Olympus Provi at 4Xmagnification used the built-in function of manual tracking in Imaris. Cells were identified as migrating spots and selected from frame to frame manually for representative migratory, oscillatory and stationary cells. Selected spots from the same cell were connected by the function of “Create Track”. Coordinate information for each track was exported from Imaris and plotted as tracks with coordinate using GraphPad Prism.

### Transmission electron microscopy

RAW264.7 cells on Mattek dishes were fixed with 2.5% glutaraldehyde in sodium cacodylate, washed in 0.1M sodium cacodylate buffer and then post-fixed in 1% osmium tetroxide with 1.5% potassium ferricyanide (ProSciTech, Thuringowa, Australia). Dishes were en-bloc stained with 1% uranyl acetate, dehydrated through a series of ethanols and capsule embedded in LX112 resin before polymerisation overnight at 60°C. Thin sections were cut from the base of snapped off capsules on a Leica Ultra Microtome and viewed on a Jeol 1011 electron microscope at 80kv (Jeol, Australasia). Images were captured using the Olympus ITEM imaging software (Soft Imaging System, Olympus, Australia).

### Zebrafish transgenic lines

Zebrafish lines were carried on a Tübingen (TU) background (Max Planck Institut für Entwicklungsbiologie, Tübingen, Germany). Compound transgenic lines were various combinations of: Tg(mpeg1:Gal4FF)^gl25^ (5), Tg(UAS-E1b:Eco:NTR-mCherry)^c264^ (54), Tg(mfap4:mTurquiose2)^xt27Tg^ (55), and Tg(mpeg1:mCherry-CAAX)^gl26^ (56). Although not utilised in these studies, the neutrophil reporter Tg(mpx:EGFP)^i114^ (57) was carried in several lines. New lines created for this study were: Tg(4xUAS:tspan4a-EGFP;cmlc2:EGFP)^gl53^, and Tg(4xUAS:scimp-EGFP;cmlc2:EGFP)^gl52^. The zebrafish experimental protocol was approved by the Monash University Ethics Committee (ERM37011) following National Health and Medical Research Council (NHMRC) guidelines. The work was approved by the Monash University Institution Biosafety Committee (PC2-N46/15). Zebrafish were maintained and bred using standard practices. Embryos were collected in E3 medium with 0.0001% w/v methylene blue and were incubated at 28°C. To prevent pigmentation, 0.003% w/v 1-phenyl-2-thiourea (PTU) (Sigma-Aldrich) was added to the E3 medium at ∼8 hrs post fertilization (hpf).

Tg(4xUAS:tspan4a-EGFP) and Tg(4xUAS:scimp-EGFP) constructs were generated by Gateway cloning (58) using destination vectors carrying Tol2 sites (59) and a cmlc2:EGFP backbone marker. Sequences and plasmids are available on request. Middle entry vectors for tspan4a-EGFP and scimp-EGFP were constructed using gBlocks (IDT). Zebrafish tspan4a sequences were as annotated in ZFIN (ZDB-GENE-040718-38) (60); of the TSPAN homologues in zebrafish, tspan4a has previously been implicated as involved in migrasome biogenesis (14). The zebrafish *scimp* gene is not annotated in ZFIN or Ensembl but has been previously recognized (20). The gBlock designs removed the tspan4 or scimp stop codons, fused the EGFP sequence in frame to their 3¢ ends, and flanked the fusion cDNAs with attB sites. Transgene constructs (25 ng/μL) were co-injected with Tol2 mRNA (50 ng/μL) into 1-cell Tg(mpeg1:Gal4FF;UAS-E1b:Eco:NTR-mCherry) embryos. Injected F0 embryos were screened for the presence of cmlc2-EGFP reporters, indicative of successful construct delivery. F0 embryos were raised to adulthood, outcrossed to identify germline founders.

### Isolation of primary zebrafish macrophages for adherent cell imaging

To isolate “germ free” macrophages from zebrafish embryos a sterilization procedure was required. Embryos were collected in E3 with 1% w/v methylene blue. After 2 hr, they were treated three times with 0.005% sodium hypochlorite (5 min) and E3 rinsing (5 min). They were then soaked in 0.01% polyvinylpyrrolidone (PVP-I; Betadine) for 30 sec, followed by two rinses with sterile E3 media and held overnight in E3 with 1X Antibiotic-Antimycotic (Gibco). Each subsequent day, dead embryos were removed and media changed with 1x Antibiotic-Antimycotic (Gibco) and 0.003% PTU. 3-5 dpf embryos were anesthetized, soaked with 0.01% polyvinylpyrrolidone (PVP-I; Betadine) (30 sec), and rinsed twice with 1X PBS. Embryos were crushed to form a cell suspension, passed through a 40 µm cell strainer, 4′,6-diamidino-2-phenylindole (DAPI, Thermo Fisher) (10 ng/µL). The FACS gating strategy was based on single cell selection, DAPI exclusion of dead cells, FSC, SCC, and mCherry expression. Sorted macrophages were cultured in DMEM (Thermo Fisher Scientific) supplemented with 10% heat-inactivated fetal bovine serum (Gibco), 2 mM GlutaMAX (Gibco), and 1X Antibiotic-Antimycotic (Gibco) on fibronectin coated slides (10 µg/mL; Sigma-Aldrich) and incubated at 28°C for 18-24 hr to permit macrophage adherence and migration. For imaging, cells were glutaraldehyde fixed (2.5%, 30 min), washed three times with 1X PBS, stained with Wheat Germ Agglutinin (WGA-Alexa Fluor 488, 1.0 µg/mL) for 10 min and washed with 1X PBS Sodium borohydride (Thermo Fisher Scientific) (1 mg/mL 7 min), aimed at reducing autofluorescence caused by fixation (61). Fixed cells were imaged using a Leica Thunder wide-field microscope.

### ZF4 cell culture, transfection and analysis

Zebrafish fibroblast ZF4 cells (CRL-2050; ATCC) (62) were cultured in DMEM/F12 (11330032; Thermo Fisher Scientific) supplemented with 10% FBS (A38401-01; Gibco), 50 μg/mL streptomycin and 50 U/mL penicillin (15070-063; Gibco) at 28°C with 5% CO_2_. For migrasome imaging, ZF4 cells were passaged at a cell density of 20,000/cm^2^ onto 35 mm FluoroDishes (FD35-100; World Precision Instrument) coated with 10 μg/mL fibronectin (F1141; Sigma-Aldrich). To simulate fibroblast cell migration, a monolayer scratch assay was used (63). Cell cultures were washed twice with phosphate-buffered saline (PBS) one day after cells were seeded on 35 mm Petri dishes. The monolayer was scratched by a cell scratcher or P200 pipette tip, incubated for 20 min, culture medium replaced, and incubated for 4-5 hr. Monolayers were washed twice with PBS prior to glutaraldehyde fixation and WGA staining as above.

To generate ZF4 cells overexpressing tspan4a, the tspan4a gBlock was PCR amplified with primers adding BamH1 and Xho1 restriction enzyme sites using Phusion® High-Fidelity PCR kit (E0553L; NEB) with a thermal cycler (T100; Bio-Rad) (98/64/72°C x35 cycles). Primers: 5’-AGTCTG**GGATCC**GGGACAAGTTTGTACAAAAAAGCAGG-3’ and 5’-CTGGATT**CTCGAG**GGGGACCACTTTGTACAAGAAAG-3’. BamH1/Xho1-digested PCR product was ligated into the pCS2+ multicloning site by standard techniques, plasmids amplified, purified and verified by Sanger sequencing (Micromon Genomics; Monash University). ZF4 cells, being lipofection-amenable (64), were transfected with Lipofectamine 3000TM transfection reagent (L3000-008; Invitrogen) at a DNA:Lipofectamine ratio of 1:1.5 and incubated for 20 hr.

ZF4 cell migrasome size was determined using Fiji/ImageJ software tools without batch-processing scripts. The manual analytical workflow steps were as detailed in (**supplementary Appendix 1**).

### Migration model, microscopy and image processing for zebrafish imaging *in vivo*

To stimulate macrophage migration *in vivo* for imaging, tails of zebrafish larvae were transected with a sterile scalpel blade as previously described (5, 41). All imaging was done on 3 dpf embryos immediately after injury up to 4 hrs.

Leica THUNDER Imager – DMi8 inverted microscope fitted with a Leica sCMOS Microscope Camera – K5 operating LAS X imaging software (Version 5.0.2). Specifications: Objectives, 40x/1.4 and 100x/1.4 oil; Image size, 2048×2048 pixels, Filter sets, FITC (Leica) (Excitation: 495 nm, Emission: 519 nm) and Texas Red (Leica) (Excitation: 596 nm, Emission: 610 nm) Zeiss LSM980 upright confocal microscope – for microscopy details, see Table 1. Zeiss LLS7 inverted lattice light sheet microscope – for microscopy details, see Table 2.

Zeiss instruments in Monash University run Zen Blue software (versions 3.8-3.9) for acquisition and initial processing. LLS7 data underwent deskewing and deconvolution (constrained iterative N=12) in Zen software. For visualization and computational analysis, Imaris software (Bitplane, versions 9.9.1 and v10.0.1) was employed. Subsequent visualization and computational analysis used ImageJ (Fiji) (v2.9.0) (65) and Imaris software. Imaris was employed for cell tracking utilizing an autoregressive motion algorithm, with manual editing of the generated tracks. Retraction fibers were counted and measured manually in Imaris.

### Quantification and Statistics

Migrating and stationary murine cells were identified by the production of RFs. Stacks of 3D fluorescence imaging of WGA-stained murine macrophages were deconvoluted in Zen Blue (Zeiss, version 3.2). Cell surfaces were constructed in Imaris with the function of “Surface” based on WGA fluorescence signals. The surfaces detail size was set as 0.144 μm. The threshold, filter of surfaces, and surface splitting and combination were adjusted for each cell to ensure the surface fitting WGA signal accurately. Surface area was measured by “Statistics” under the function of “Surface” and plotted in GraphPad Prism 10 (**Fig 8C**). ROI of each cell was manually selected based on WGA-stained cell membrane. Integrated TNF fluorescence intensities were measured for each ROI and plotted in GraphPad Prism 10 (**Fig 8D**). One data point represents one reading from a cell.

The novel parameter RF AUC/cell/min (**Fig 2G**), a surrogate proportional to retraction fiber incidence, was computed by determining the area under the curve from a plot of retraction fiber number at each frame for the duration of the movie for the cells in **Fig 2E-F** and dividing by the length of the movie, thereby weighting each cell equally regardless of movie length. Figures were constructed in Adobe Illustrator (2022, 2023).

For all studies, descriptive and analytical statistics were prepared in GraphPad Prism (v10.0.0, GraphPad Software, Boston, Massachusetts USA). The figure legends include details on the statistical tests performed, n values, and *p* values. All tests are two-tailed; *p* values < 0.05 (adjusting where necessary for multiple comparisons) are interpreted as significant.

## ACKNOWLEDGEMENTS

We thank Monash AquaCore staff for technical assistance with fish husbandry, Dr S. Firth and Monash Micro Imaging staff for assistance with zebrafish imaging, and Prof. P. Currie for encouragement. We thank Tatiana Khromykh for expert assistance with murine cell culture and acknowledge all members of the Stow lab for help and advice. Cell imaging and image processing for murine cells was performed in IMB Microscopy.

## Funding

Parts of this work were supported by the Australian Research Council (DP230100504 to JLS, GJL, RZM, NDC; DP210103263 and LE220100138 to GJL and the ARC Centre of Excellence in Quantum Biotechnology CE230100021 to JLS, by the National Health and Medical Research Council of Australia (APP1176209 to JLS) and by Chan Zuckerberg Initiative (2023-329684 to NDC). Graduate scholarships were funded by Monash University (to A. Anbarlou and International Tuition Scholarship), and by Queensland University of Technology (to A. Ashraf) and by The University of Queensland (to WW) and by UQ’s Institute for Molecular Bioscience (Global Challenges scholarship to HS). The Australian Regenerative Medicine Institute is supported by grants from the State Government of Victoria and the Australian Government.

## Abbreviations

BMM: bone borrow-derived macrophage
CSF-1: colony stimulating factor 1
EGFP: enhanced green fluorescent protein
FN: fibronectin
HMDM: human monocyte-derived macrophages
LLSM: lattice light sheet microscopy
LPS: lipopolysaccharide
LUT: lookup table
MIP: maximum intensity projection
RF: retraction fiber
TNF: tumor necrosis factor
TAPI: TNF alpha processing inhibitor
TEM: tetraspanin-enriched microdomain
WGA: wheat germ agglutinin
pTRAP: palmitoylated transmembrane adaptor protein

## Supporting information

**Supplementary figure 1.**
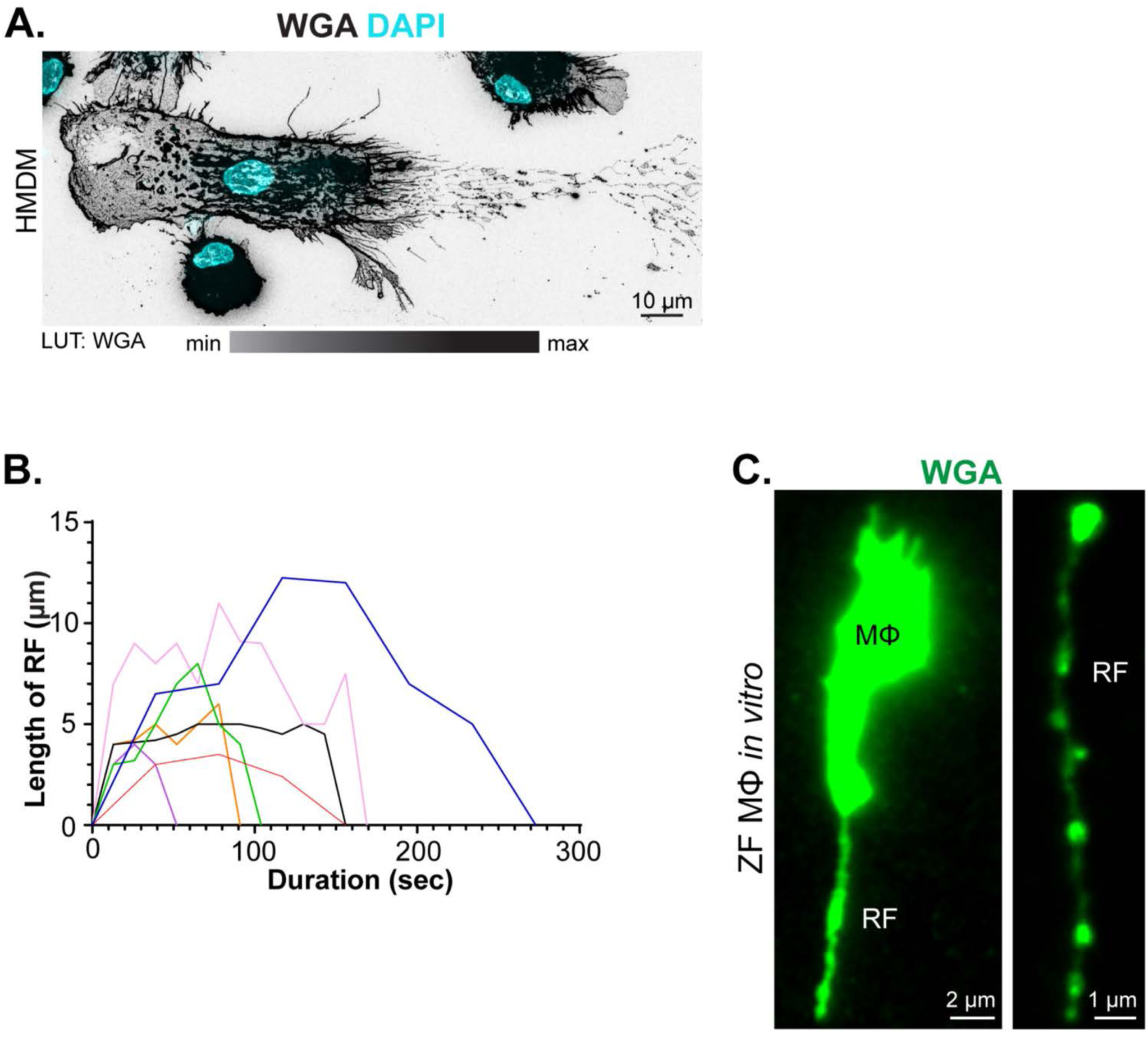
Retraction fibers and migrasomes of migrating macrophages *in vitro* and persistence *in vivo*. **A.** HMDMs differentiated under CSF-1 treatment, re-plated on FN-coated coverslips and treated with IL-4 for 12 hrs. Migrating macrophages were fixed and stained with Alexa Flour 488-WGA (LUT: inverted Grays) and DAPI (inverted Cyan). **B.** Length and persistence of RFs (n=7). RF and migrasome formation by zebrafish macrophages *in vivo* in Tg(mfap4:mTurquoise2;mpeg1:mCherry-CAAX) macrophages. **C.** RF and migrasome formation by zebrafish macrophages, demonstrated *in vitro*. FACS-purified zebrafish macrophages were plated on FN-coated plates, fixed, and stained with Alexa Flour 488-WGA. Left image shows macrophage soma (Mf) and attached single RF. Right image shows a detached WGA-stained fragment elsewhere in the preparation, appearances consistent with a RF carrying migrasomes. Leica Thunder widefield images. **A** and **C**, Scalebars as indicated.

**Supplementary figure 2.**
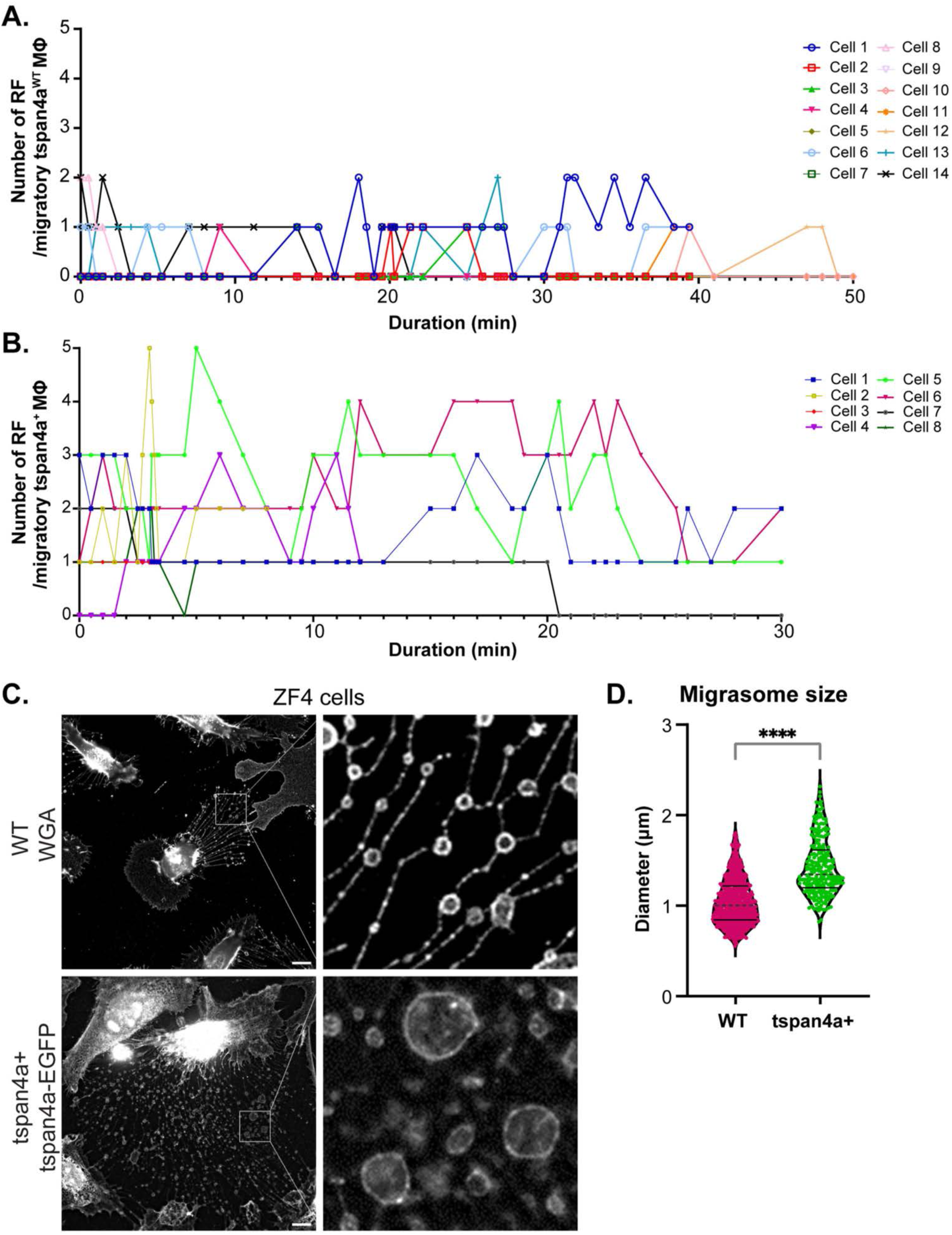
Impact of tspan4a+ on retraction fiber and migrasome formation in zebrafish macrophages *in vivo* and ZF4 fibroblasts *in vitro*. **A-B.** RF number for tspan4a+ (n=8) and WT (n=14) macrophages. WT macrophages displayed £ 2 RFs (**A**) whereas tspan4a+ displayed 1-5 RFs (**B**) for most of the time. Data for **A-B** from MIP images, Zeiss LSM980 confocal microscope. **C-D.** Migrasome formation by ZF4 fibroblast cells *in vitro*. **C.** Upper row panels show unmanipulated ZF4 cells, fixed and stained with Alexa 488-WGA. Lower row panels show ZF4 cells transduced to express tspan4a-EGFP, live imaged. Insets on right are magnified views of areas as shown, highlighting retraction fibers and their associated migrasomes. Scalebars as indicated. **D.** The diameter of migrasomes formed by tspan4a overexpressing ZF4 cells is increased by 37%. The number of randomly-selected migrasomes for each ZF4 genotype: WT n = 834, tspan4a+ n = 240, each collected from random fields of multiple cells. Data are from MIP images, Leica Thunder microscope. Data are presented with median and quartiles. Mann-Whitney test. **** *p*<0.0001.

**Supplementary figure 3.**
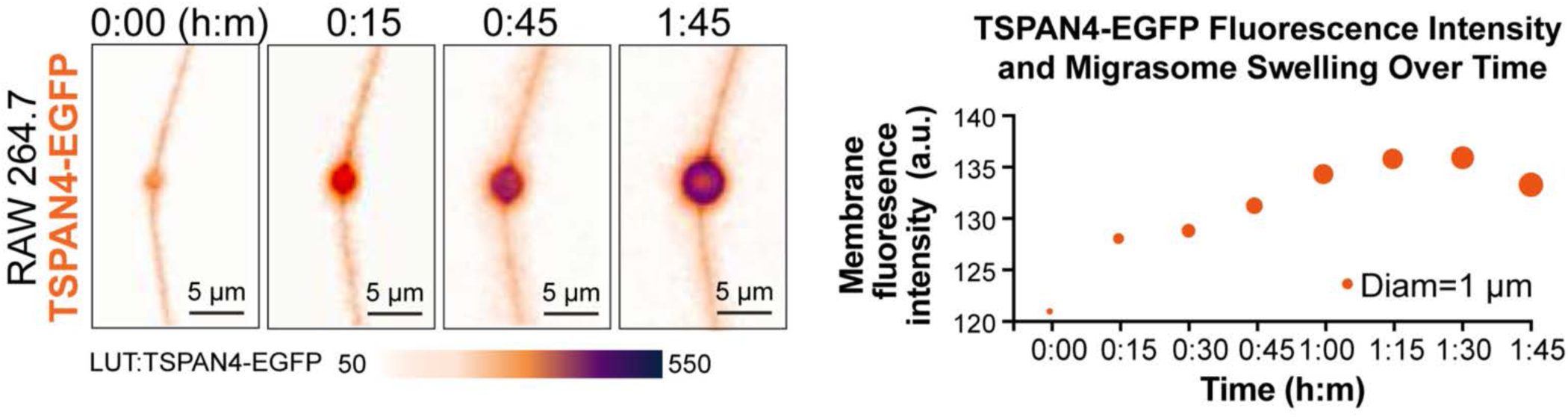
TSPAN4-EGPF packing in murine macrophage migrasome expansion. Left panel: A migrasome expansion recorded in sequential frames from live imaging in RAW264.7 TSPAN4-EGFP expressing macrophages. Vesicle expansion and TSAPN4-EGFP accumulation during migrasome formation is evident. LUT: inverted gem for TSPAN4-EGFP. Scalebars as indicated. Right panel: For this migrasome in left panel, corresponding plot of temporal changes in migrasome diameter and TSPAN4-EGFP fluorescence intensity of the migrasome membrane. The dot size represents migrasome diameter at the time point showing in x axis.

**Supplementary figure 4.**
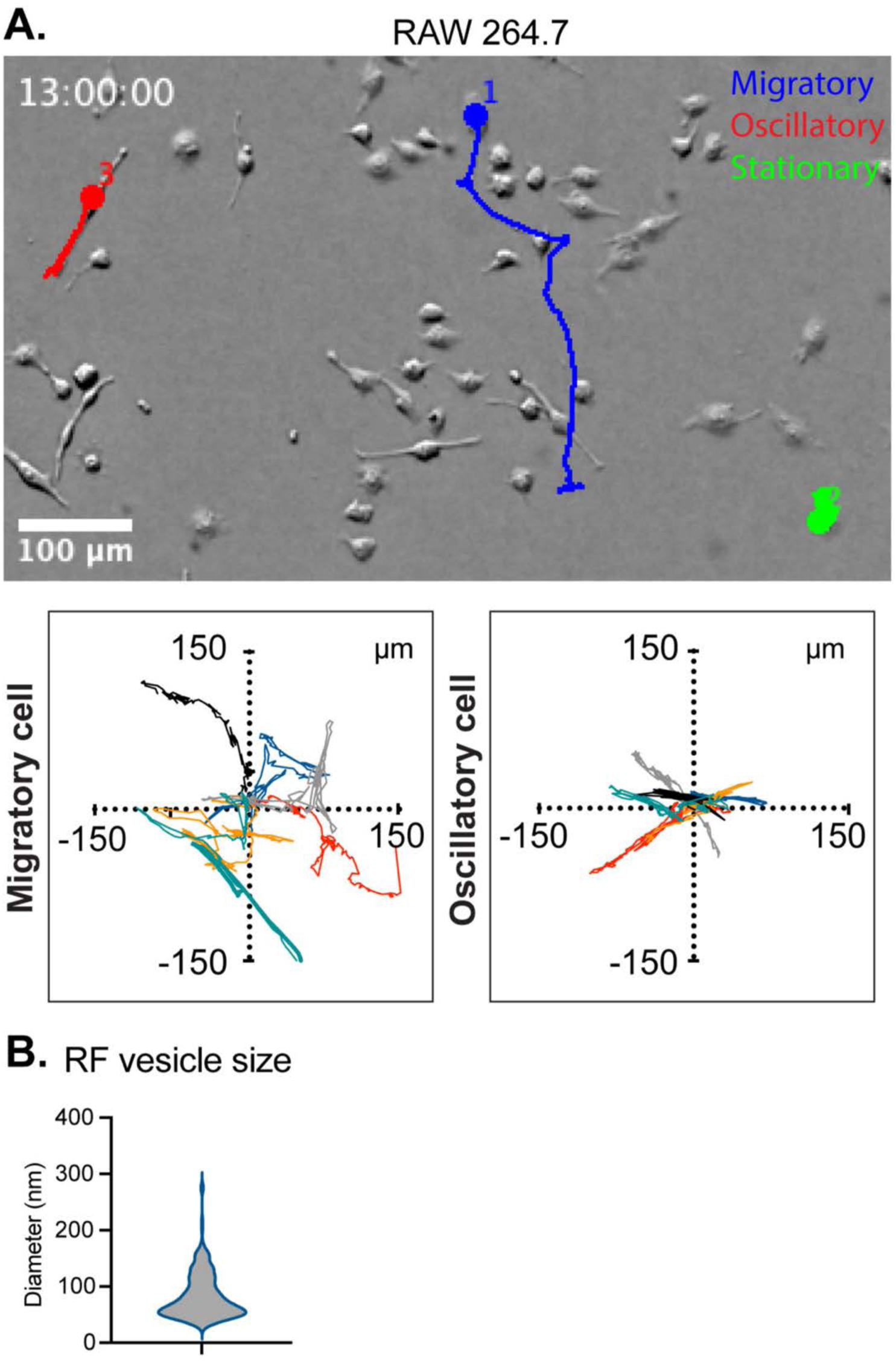
Tracks of migratory and oscillatory macrophages *in vitro* and RF-derived extracellular vesicles. **A.** Migratory (blue), oscillatory (red) and stationary (green) trajectories of macrophages in a low-magnification field of view (top). Migration trajectories from migratory cells (bottom left) and oscillatory cells (bottom right). Bright field live imaging of RAW 267.4 cells in CSF-1 recorded with 4X objective for > 16 hrs. Cells mapped as spots in Imaris and manually tracked using built-in tracking algorithm. Time series data of coordinates plotted by Prism GraphPad. See **Video 4**. Scalebars as indicated. **B.** Violin graph shows diameters of vesicles on matrix. Diameter was measured in FIJI based on EM dataset. n = 223 vesicles.

**Supplementary figure 5.**
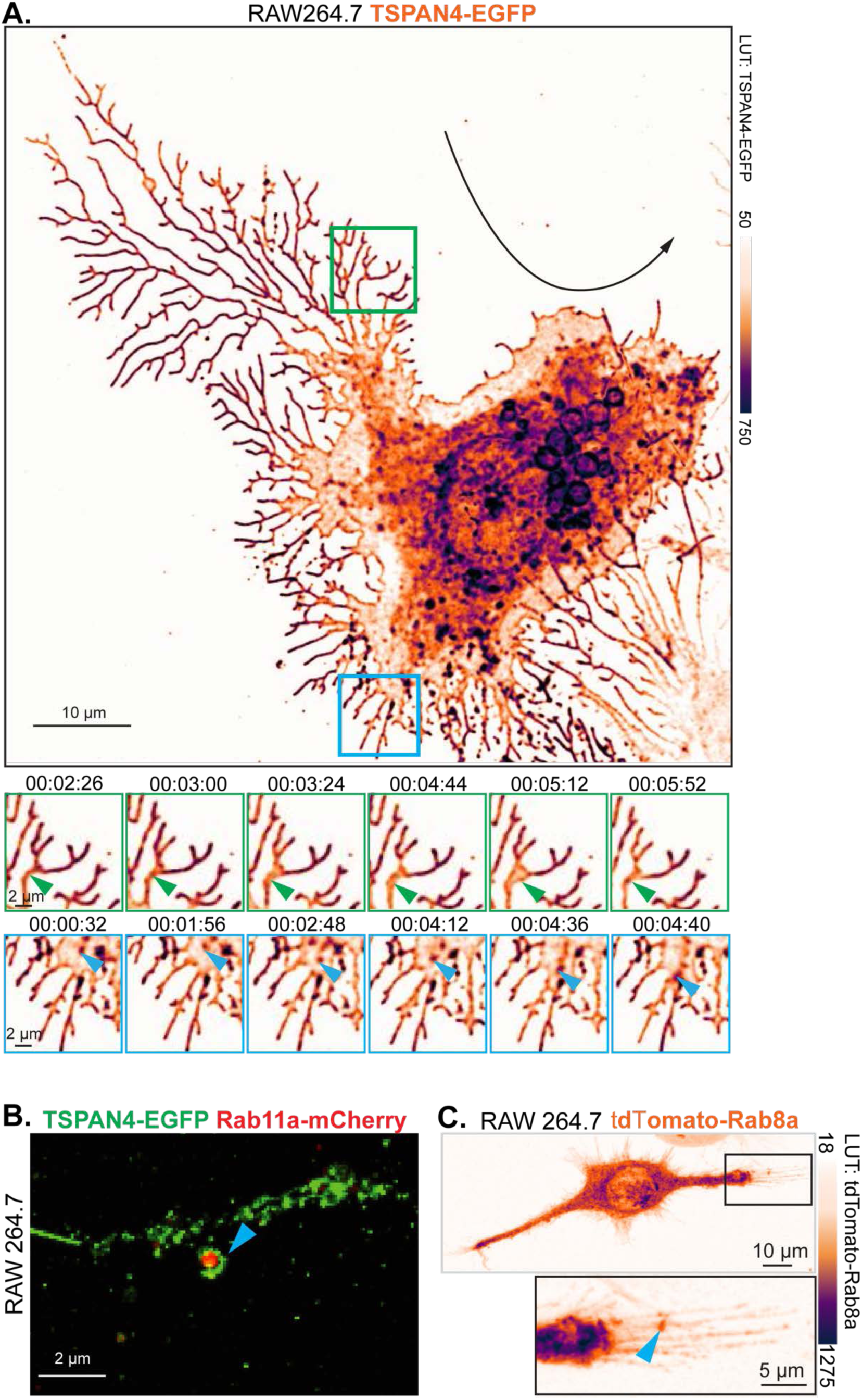
Cytoplasm and vesicle trafficking at the base of macrophage RFs. **A.** RAW264.7 macrophages treated with CSF-1 for 6 hrs. Live cells were imaged by LLSM at 4 sec/frame. The green inset shows cytoplasm being pushed into RF bases (green arrows). The blue inset shows TSPAN4-EGFP-positive vesicles traveling into and out of RF bases (blue arrowheads). LUT: inverted gem for TSPAN4-EGFP. See **Video 8**. **B.** RAW264.7 cells co-transfected with TSPAN4-EGFP (green) and mCherry-Rab11a (red) on FN-coated MatTek dishes for live imaging. mCherry-Rab11a was detected in migrasomes (blue arrowhead). **C.** CSF-1 treated WT RAW264.7 cells transfected with tdTomato-Rab8a. Cells were incubated in CSF-1 overnight post transfection, fixed and imaged on a wide field fluorescence microscope. LUT: inverted gem for Tdtomato-Rab8a. **A-C**, scalebars as indicated.

**Supplementary figure 6.**
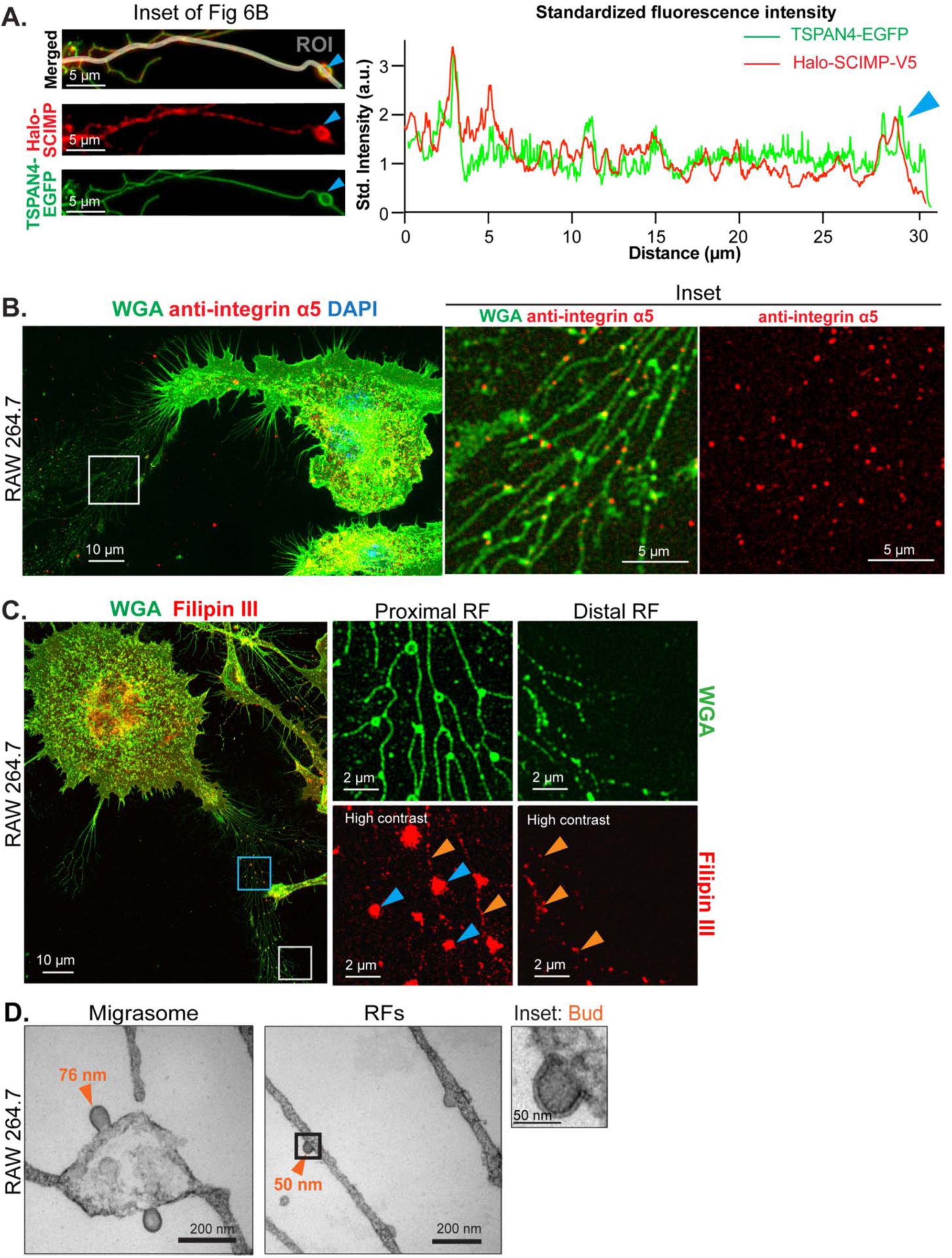
TEM composition in murine macrophage RFs and migrasomes. **A.** TSPAN4-EGFP (green) and Halo-SCIMP (JF549 HaloTag^®^ ligand, red) colocalization analysis. ROI was selected along the RF in an inset of a RF with a migrasome from **Fig 6B**. Fluoresence intensity of each channel was normalized as standard fluorescence intensity (Std. intensity). Std. intensity for both channels was plotted using GraphPad Prism, with peaks indicating protein enrichment at specific positions. **B.** Immunostaining of integrin ⍺5 (CD49e, staining with anti-CD49e primary antibody followed by a secondary antibody with Alexa Fluor 594, red) in migrating macrophages, showing localization in RF and migrasomes (insets). Alexa Fluor 488-WGA (green) and DAPI (blue) label membranes and nuclei respectively. **C.** Filipin III (LUT: Red) staining for cholesterol detection in migrating macrophages, showing localization in RF and migrasomes. Alexa Fluor 488-WGA (green) was applied to stain cell membranes. The blue-boardered inset of a proximal RF shows cholesterol enrichment in migrasomes (blue arrowheads) and localization along RFs (orange arrowheads) with contrast enhancement. The white-boardered inset of a distal RF under enhanced contrast highlights cholesterol (orange arrowheads) along RFs. **D.** CSF-1 treated RAW264.7 cells, fixed and processed as thin sections for transmission EM. Images show RFs and migrasomes budding. **A-D**, scalebars as indicated.

**Supplementary figure 7.**
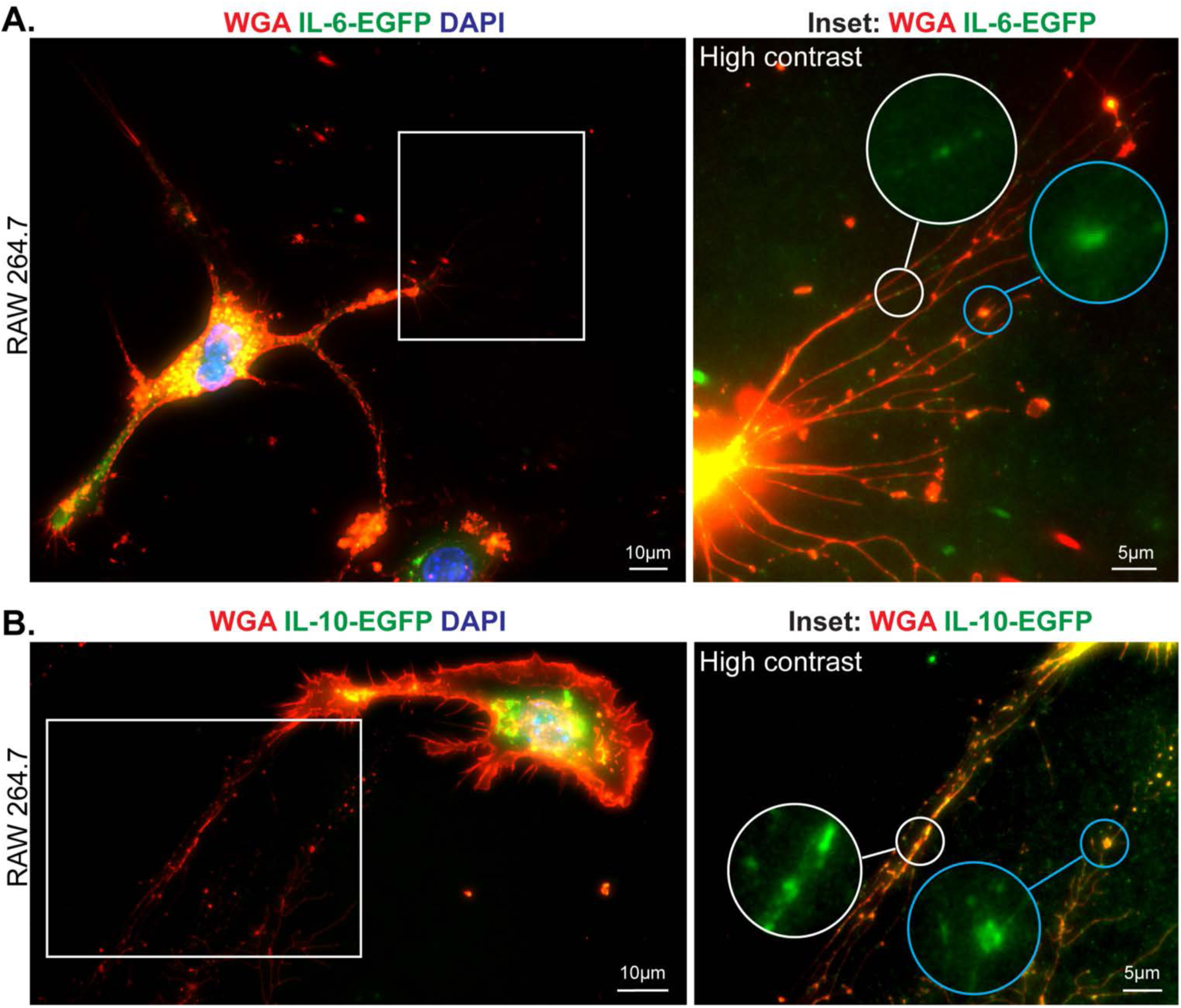
Soluble and membrane-bound cytokines traffic into RFs and migrasomes. **A** and **B**: RAW264.7 cells transfected with IL-6-EGFP (**A**) or IL-10-EGFP (**B**) by electroporation. After overnight recovery, cells were replated on FN-coated coverslips and treated with CSF-1 for 16 hrs, then fixed and stained with Alexa Fluor 647-WGA (red) and DAPI (blue). Contrast enhanced insets show the accumulation of IL-6-EGFP (green) (**A**) and IL-10-EGFP (green) (**B**) along RFs (white border insets and migrasomes (blue border insets). Insets are presented as high contrast and only EGFP channel is shown. Scalebars as indicated.

**Supplementary figure 8.**
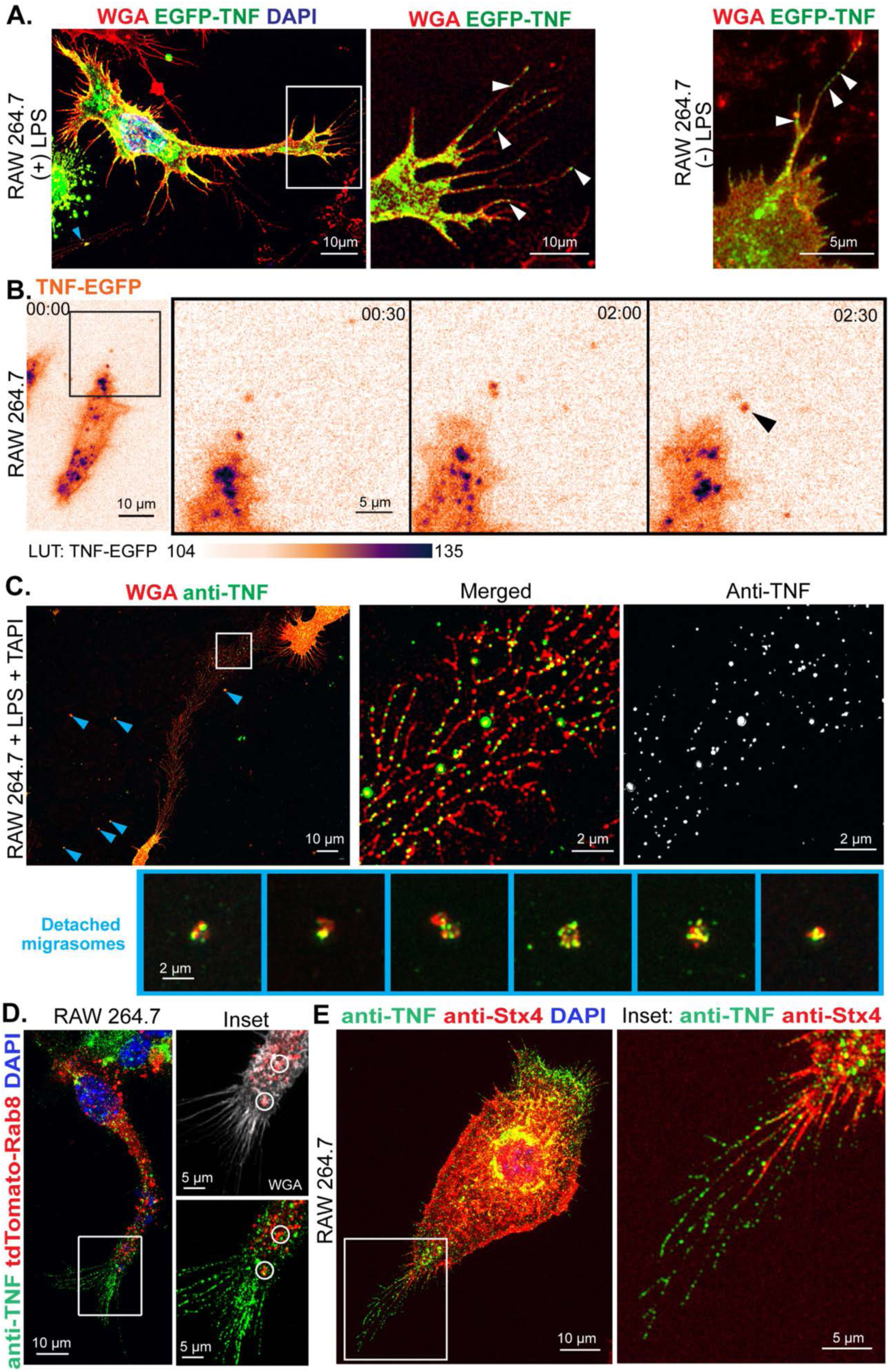
Recombinant or endogenous TNF trafficking and presentation along RFs and in migrasomes. **A.** RAW264.7 cells treated with CSF-1 overnight, then transfected with EGFP-TNF (green). After a 2-hrs recovery, cells were either stimulated with LPS for 4 hrs before fixation (left panel) or fixed without LPS stimulation (right panel). White arrows indicate EGFP-TNF distribution along RFs. **B**. RAW264.7 cells were transfected with TNF-EGFP. Live imaging was performed 2 hrs after transfection on wide field fluorescence microscope with a scanning interval of 30 sec. Insets and black arrows show translocation of a TNF granule into migrasomes (black arrowheads). LUT: inverted gem for TNF-EGFP. **C.** Endogenous TNF presentation in RFs and migrasomes. RAW264.7 cells were cultured on FN-coated substrates and incubated with CSF-1 for 16 hrs, followed by a 2-hrs LPS and TAPI incubation before fixation. Fixed cells were immunostained with TNF antibody, followed by an Alexa Fluor 488 conjugated secondary antibody (green) along with Alexa Fluor 647-WGA (red) and DAPI (blue) staining. Upper panels: pro-TNF (green) retained on FN matrix were imaged together with RFs and detached migrasomes (WGA-stained vesicles). Lower panels: TNF-contained migrasomes detached from RF while remained on FN matrix. **D.** tdTomato-Rab8-transfected RAW264.7 cells were treated with LPS and TAPI simultaneously for 2 hrs. Cells were then fixed, permeabilized, immunostained for TNF, and stained with WGA and DAPI for membrane and nuclei. Insets show the rear end of the cell with RFs. Grays LUT was used for anti-TNF in the top inset, and co-localization of tdTomato-Rab8a and TNF is circled. **E.** Cells were treated with LPS and TAPI for 2 hrs and then fixed and immunostained with TNF antibody (secondary antibody conjugated with Alexa Fluor 488, green) and Stx4 antibody (secondary antibody conjugated with Alexa Fluor 594, red) simultaneously. Cell nuclei were stained with DAPI (blue). **A-E**, Scalebars as indicated.

**Supplementary figure 9.**
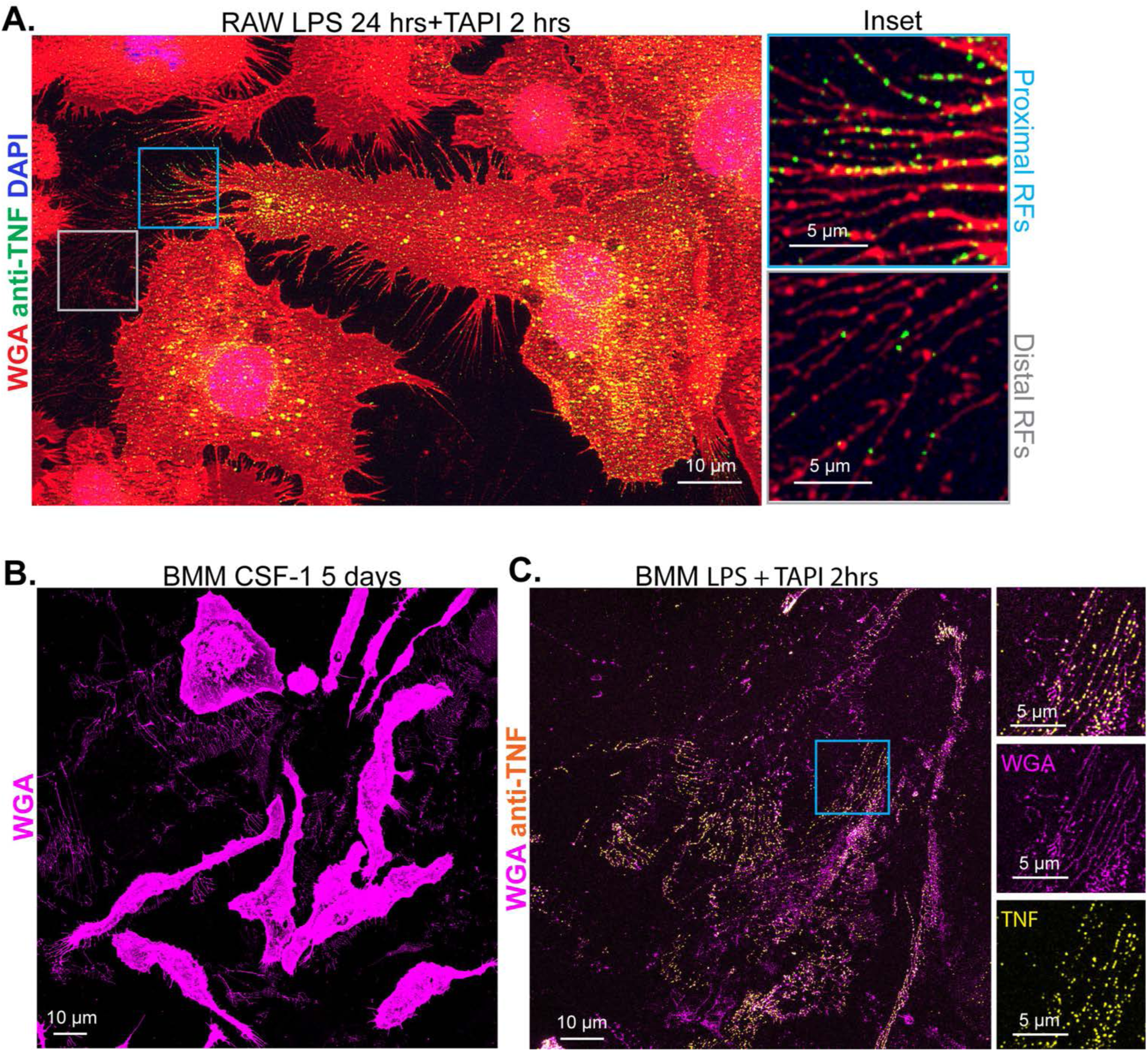
TNF retention in macrophage RF trails and on the matrix. **A.** RAW264.7 cells in LPS for 24 hrs, with TAPI added for last 2 hrs. Insets show endogenous TNF staining (shown with Alexa Fluor 488-conjugated secondary antibody, green) in proximal RFs (blue box) and distal RFs (white box). Alexa Fluor 647-WGA (red) and DAPI (blue) were applied to visualize cell membrane and nuclei. **B.** Macrophages were differentiated from bone marrow monocytes over 5 days in the presence of CSF-1, showing the formation of dense RFs on the matrix. On the 5^th^ day of differentiation, cells were fixed and stained with Alexa Fluor 488-WGA (LUT: Magenta). **C.** BMMs were pretreated with CSF-1 for 16 hrs after replating, followed by stimulation with LPS and TAPI for 2 hrs. Cells were then fixed and immunostained using an anti-TNF antibody and an Alexa Fluor 488-conjugated secondary antibody (LUT: Yellow). Alexa Fluor 647-WGA (LUT: Magenta) was used to visualize RFs. Dense RF webs colocalized with pro-TNF were observed in certain fields of view on coverslips. **A-C**, scalebars as indicated.

## VIDEO LEGENDS

**Video 1. Recruitment of zebrafish macrophages into injury site *in vivo*.**

Source video for figure 1B, showing Tg(mpeg1:Gal4FF;UAS:NTR-mCherry) zebrafish macrophages leaving the circulation and migrating towards the tail injury site. Zeiss LSM980 confocal images. Time stamp = hh:mm:ss.ms. Scalebar = 50 μm.

**Video 2. Retraction fiber and migrasome formation by zebrafish macrophage *in vivo*.** Source video for figure 1H, showing a migrating Tg(mpeg1:Gal4FF;UAS:NTR-mCherry) zebrafish macrophage, forming a retraction fiber which fragments. See embedded video labels. Zeiss LSM980 confocal images. Time stamp = hh:mm:ss.ms. Sequence 1: MIP images, mCherry in black; followed by Sequence 2: surface rendered, mCherry in white/grey. Scalebar = 10 μm.

**Video 3. Retraction fiber and migrasome formation by tspan4a-overexpressing zebrafish macrophage *in vivo*.**

Source video for figure 2B, showing a migrating Tg(mpeg1:Gal4FF;UAS:NTR-mCherry;UAS:tspan4a-EGFP) zebrafish macrophage. Note multiple retraction fibers fragment, each showing strong tspan4a-EGFP (yellow) signal. See embedded video labels. A follower macrophage appearing top right at t=∼17:00 has a lower level of tspan4a:EGFP expression. Zeiss LSM980 confocal MIP images. Time stamp = hh:mm:ss.ms. mCherry cytoplasm (magenta), tspan4a-EGFP reporter (yellow), merged/overlay (white). Scalebar = 5 μm.

**Video 4. Macrophage migration trajectories *in vitro*.**

Source video for **supplementary figure 4A**, showing RAW 264.7 macrophages randomly migrating in the presence of CSF-1. Their trajectories were classified into three types: Migratory (blue), Oscillatory (red), and Stationary (green). Live imaging of macrophages was recorded for 44 hrs (partially presented in this video) with an interval of 15 mins. Trajectories were depicted using the ‘Manual Tracking’ function in FIJI. Time stamp: hh:mm:ss. Scalebar = 100 μm.

**Video 5. RFs and migrasomes formed by migratory macrophages *in vitro***

Source video for figure 4A. RAW 264.7 cells stably expressing TSPAN4-EGFP were cultured on FN-coated MatTek dishes and imaged using widefield fluorescence microscopy in the presence of CSF-1 for 16 hrs (timeline cropped according to cell behaviors) at 15 mins per frame. Raw data were cropped in Imaris and deconvoluted with Microvolution in FIJI, as described in Methods and Materials. A MIP was presented, and an inverted gem LUT was applied for TSPAN4-EGFP. Time stamp: hh:mm:ss. Scalebar = 10 μm.

**Video 6. RFs and migrasomes formed by oscillatory macrophages *in vitro***

Source video for figure 4A. RAW 264.7 cells were treated and imaged the same as described for **Video 5**. An oscillatory cell produces RFs and migrasomes while switching migrating directions several times. Time stamp: hh:mm:ss. Scalebar = 10 μm.

**Video 7. Retraction fiber and migrasome maturation from a tspan4a-overexpressing zebrafish macrophage *in vivo*.**

Source video for figure 4D, showing a migrating Tg(mpeg1:Gal4FF;UAS:NTR-mCherry;UAS:tspan4a-EGFP) zebrafish macrophage. The retraction fiber breaks, the migrasomes along its length enlarge, dissociate and disperse. See embedded video labels. Zeiss LSM980 confocal MIP images. Time stamp = hh:mm:ss. mCherry cytoplasm (black) channel only. Scalebar = 5 μm.

**Video 8. Cytoplasm and vesicle trafficking at the base of RFs in migrating macrophages *in vitro***

Source video for **supplementary figure 5A**. RAW 264.7 macrophages expressing TSPAN4-EGFP were treated with CSF-1 for 6 hrs. Live imaging was performed on LLSM at 5 sec/frame. MIP was presented and inverted gem LUT was applied for TSPAN4-EGFP. Time stamp: hh:mm:ss. Scalebar = 10 μm.

**Video 9. Aggregation of recycling endosomes at the bases of processes form retraction fibers in migrating macrophages *in vivo*.**

Source video for figure 5B, showing a migrating Tg(mpeg1:mCherry-CAAX) zebrafish macrophage. This elongated branched macrophage displays multiple freely mobile internal membrane-labelled endosomes. As retraction fibers form on several protrusions, accumulation of these membrane-labelled endosomes at the base of retraction-fiber forming cellular processes is evident, both as individual endosomes and also as aggregation of the membrane-localized reporter. See embedded video labels. Zeiss LLS7 lattice light sheet MIP images. Time stamp = hh:mm:ss.ms. mCherry cytoplasm channel only. Scalebar = 5 μm.

---

**Supplementary Appendix 1. Image processing for *in vitro* zebrafish cell studies** ZF4 cell migrasome size was determined using Fiji/ImageJ software tools without batch-processing scripts. The manual analytical workflow steps were as follows:

(1) Imports an image (raw data) into Fiji/ImageJ;
(2) Sets the “Unit of length” to micron and modifies the “Distance in pixels” to the desired value in the “Set Scale” panel;
(3) Enhances the image contrast by adjusting values in the “Brightness/Contrast” panel;
(4) Applies the “Freehand Line” tool to draw and label retraction fiber;
(4) Adds selected retraction fibers to the “ROI Manager”;
(5) Classifies the parental, branch and inter-migrasome retraction fibers based on their length by marking the retraction fibers in “ROI Manager”;
(7) Defines the half-point of the parental retraction fibers and delimits proximal and distal regions in accordance with the half-point of the parental retraction fibers;
(8) Applies the “Freehand” tool to draw and label migrasomes. Only the migrasomes with lumen were selected;
(9) Adds selected migrasomes to the “ROI Manager”;
(10) Classifies the proximal, distal, tip and nodal migrasomes according to their location by marking ROIs (region of interest) in “ROI Manager”;
(11) Uses the “Measure” function to get the measurement results of retraction fibers and migrasomes;
(12) Save the measurement results in CSV format for further statistical analysis.

**S1. Table.** Zeiss LSM980 confocal microscopy conditions.

| Reporter line | Bit depth | Objective mag./NA | Scaling $\mu\text{m}$ | Scan mode sec | Pinhole $\mu\text{m}$ | Channel | | | | |
| --- | --- | --- | --- | --- | --- | --- | --- | --- | --- | --- |
|  |  |  |  |  |  | No. | Laser nm | Laser % | Master gain | Filter nm |
| Tg(mfap4:mTurquoise2;<br>mpeg1:mCherry-CAAX) | 16 | 20X/1.0 | x: 0.354-0.442<br>y: 0.354-0.442<br>z: 0.420 | Frame 16-36 | 28 | 1 | 405 | 1.5 | 750 | 411-570 |
|  |  |  |  |  |  | 2 | 561 | 2.0 | 750 | 578-696 |
| Tg(mpeg1:Gal4FF;<br>UAS:NTR-mCherry) | 16 | 20X/1.0 | x: 0.276-0.414<br>y: 0.276-0.414<br>z: 0.560 | Frame 14-26 | 37 | 1 | 561 | 1.0 | 750 | 578-696 |
| Tg(mpeg1:Gal4FF;<br>UAS:NTR-mCherry;<br>UAS:tspan4a-EGFP) | 16 | 20X/1.0 | x: 0.276<br>y: 0.276<br>z: 0.480-1.000 | Frame 17-36 | 31 | 1 | 488 | 3.0 | 750 | 490-579 |
|  |  |  |  |  |  | 2 | 561 | 1 | 750 | 578-696 |
| Tg(mpeg1:Gal4FF;<br>UAS:NTR-mCherry;<br>UAS:scimp-EGFP) | 8 | 20X/1.0 | x: 0.244<br>y: 0.244<br>z: 1.000 | Frame 19 | 31 | 1 | 488 | 7.5 | 850 | 490-579 |
|  |  |  |  |  |  | 2 | 561 | 4.5 | 850 | 578-696 |

**S2. Table.** Zeiss LLS7 lattice light sheet microscopy conditions.

| Reporter line | Bit depth | Objective mag./NA | Size of light sheet | Scaling $\mu\text{m}$ | Exposure time msec<br>frame sec | Channel | | | |
| --- | --- | --- | --- | --- | --- | --- | --- | --- | --- |
|  |  |  |  |  |  | No. | Laser nm | Laser % | Filter nm |
| Tg(mpeg1:mCherry-CAAX) | 16 | 40X/1.0 | Sinc3<br>30x1000 | x: 0.145<br>y: 0.145<br>z: 0.145 | 10<br>7.5 | 1 | 561 | 10 | 576-617 |

## Notes

### Competing Interest Statement

The authors have declared no competing interest.

